# A Role for the Locus Coeruleus in the Modulation of Feeding

**DOI:** 10.1101/2019.12.18.881599

**Authors:** Natale R. Sciolino, Madeline Hsiang, Christopher M. Mazzone, Leslie R. Wilson, Nicholas W. Plummer, Jaisal Amin, Kathleen G. Smith, Christopher A. McGee, Sydney A. Fry, Cindy X. Yang, Jeanne M. Powell, Michael R. Bruchas, Alexxai V. Kravitz, Jesse D. Cushman, Michael J. Krashes, Guohong Cui, Patricia Jensen

## Abstract

Recent data suggest that LC-NE neurons play a role in fear-induced suppression of feeding, but their endogenous activity in naturally behaving animals has not been explored. We found that endogenous activity of LC-NE neurons was enhanced during food approach and suppressed during food consumption, and that these food-evoked LC-NE responses were attenuated in sated mice. Interestingly, visual-evoked LC-NE activity was also attenuated in sated mice, demonstrating that internal satiety state modulates LC-NE encoding of multiple behavioral states. We also found that food intake could be attenuated by brief or longer durations of LC-NE activation. Lastly, we demonstrated that activation of LC neurons suppresses feeding and enhances avoidance and anxiety-like responding through a projection to the lateral hypothalamus. Collectively, our data suggest that LC-NE neurons modulate feeding by integrating both external cues (e.g., anxiogenic environmental cues) and internal drives (e.g., nutritional state).

## INTRODUCTION

Noradrenergic neurons of the locus coeruleus (LC-NE) are well-known for modulating high-arousal behavioral states such as orienting, waking and startle*(1-6)*. In contrast, much less is known about their contribution to feeding. To date, investigation into the role of central NE in feeding have primarily focused on noradrenergic neurons in the nucleus of the solitary tract (NTS), demonstrating that these cells can promote or inhibit feeding*(7-9)* through projection-specific pathways to the arcuate*(10)* or parabrachial nucleus (PBN)*(11)*. However, an early *in vivo* electrophysiology study in rats noted inhibition of LC neurons during sucrose consumption*(12)* and a recent study in mice reported chemogenetic inhibition of LC-NE neurons prevented fear-induced suppression of feeding*(13)*. While these data suggest that LC-NE neurons may play a role in the modulation of feeding, their endogenous activity in naturally behaving animals has not been explored. To address this question, we combined behavioral and metabolic approaches with fiber photometry calcium imaging, chemogenetics, and optogenetics. We found that endogenous activity of LC-NE neurons was enhanced during food approach and suppressed during food consumption, and that these food-evoked LC-NE responses were attenuated in sated mice. Interestingly, visual-evoked LC-NE activity was also attenuated in sated mice, demonstrating that internal satiety state modulates LC-NE encoding of multiple behavioral states. We also found that activation of LC-NE neurons resulted in the suppression of feeding when stimulation was paired to feeding in a temporally selective and behavioral relevant manner. Lastly, we demonstrated that activation of LC-NE neurons results in the suppression of feeding through a projection to the lateral hypothalamus.

## RESULTS

### LC-NE Activity Is Enhanced During Food Approach and Suppressed During Feeding in a Manner Influenced by Nutritional State

To identify the natural activity patterns of LC-NE neurons during feeding, we used fiber photometry*(14, 15)* to monitor fluorescent calcium activity using GCaMP6f in LC-NE neurons during food approach and consumption **(Fig. 1A)**. Flp-dependent GCaMP6f and tdTomato (tdT) AAVs were bilaterally injected into the LC (LC^GCaMP/tdT^) of mice expressing Flpo recombinase from the endogenous noradrenergic-specific dopamine-beta hydroxylase locus *(Dbh*^*Flpo*^*)(16)* **(Fig. 1A)**. To minimize the effects of stress and novelty during the experiments, LC^GCaMP/tdT^ mice were habituated for several days to eat chow pellets from the feeding experimentation device (FED)*(17)*. To determine if hunger state modulates LC-NE neuronal responses to food, mice were fasted overnight, and unilateral LC responses were recorded as mice fed to satiety. Early in the session, we found LC-NE neurons were activated during food approach and inhibited during consumption of pellets **(Fig. 1B-D)**, demonstrating a divergent pattern of LC activity during appetitive and consummatory behaviors. The approach-related LC-NE neuronal responses were completely abolished later in the session when mice had consumed more food, whereas the consummatory-related neuronal responses were slightly attenuated **(Fig. 1B-D)**. To determine if transition from hunger to satiety had a gradual impact on food-related LC activity across the session, we next performed linear regression analyses. We found that LC-NE neuronal responses during approach and consumption were gradually attenuated with successive pellets in the session **(Fig. 1E)**. Importantly, velocity remained stable as satiation developed across the session **(Fig. 1F**), indicating that ambulation did not drive satiety-related changes in LC-NE activity during feeding. Taken together, our findings demonstrate that endogenous activity of LC-NE neurons is enhanced during food approach and suppressed during food consumption, and that these food-evoked responses are attenuated by satiety.

**Fig. 1.**
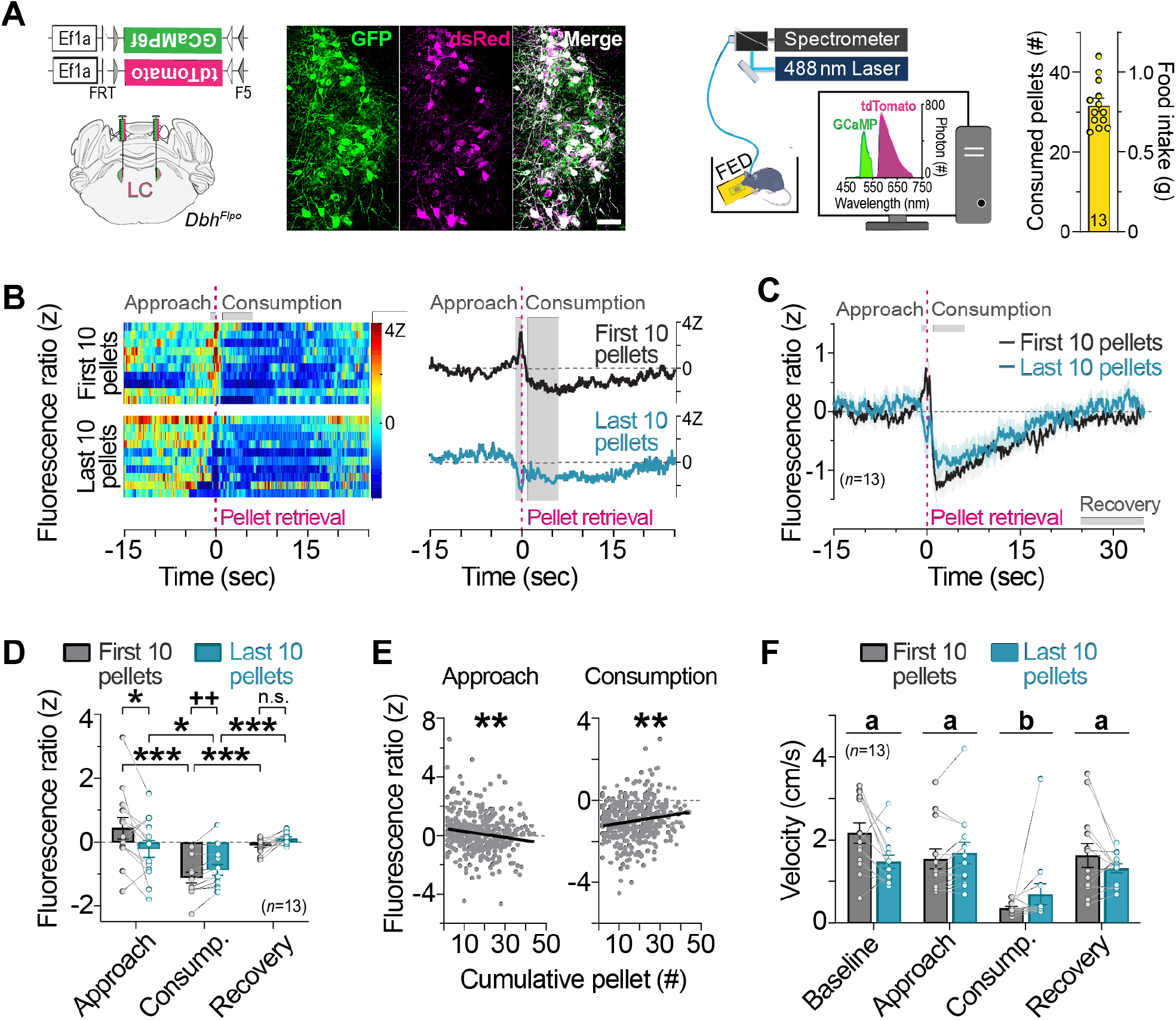
LC-NE activity is increased during food approach and suppressed during feeding in a manner influenced by nutritional state. **(A)** *Left*. Flp-dependent viral genetic strategy for co-expression of GCaMP6f and tdTomato (tdT) in LC-NE neurons. *Middle*. Coronal view of the locus coeruleus from an LC^GCaMP/tdT^ mouse immunostained for GCaMP6f (GFP antibody) and tdT (dsRed antibody). Scale, 50-µm. *Right*. Schematic of *in vivo* fiber photometry setup and feeding experimentation device (FED). Average food intake across the 1-hr session. Data are mean ± SEM. *n*=13 LC^GCaMP/tdT^ mice. **(B)** Fluorescence ratio expressed as a z-score aligned to pellet retrieval during the first ten *(top)* and last ten pellets *(bottom)* of the session from a representative fasted LC^GCaMP/tdT^ mouse. Single trials are represented in the heatmap *(left)*, and average z-score fluorescence ratio is represented in the trace *(right)*. **(C)** Average z-score fluorescence ratio aligned to pellet retrieval in fasted LC^GCaMP/tdT^ mice during the first ten pellets and last ten pellets of the session. Data are mean ± SEM. *n*=13 LC^GCaMP/tdT^ mice. **(D)** Average z-score fluorescence ratio during feeding-related behaviors. Two-way repeated measures ANOVA, nutritional state x behavior interaction: *F*_2,24_=8.819, *P*=0.0013. Bonferroni post-hoc test, \*\*\**P*<0.001, \**P*<0.05. n.s., non-significant. Paired samples *t*-test: *t*_12_=3.807, ^++^*P*<0.01. Data are mean ± SEM. *n*=13 LC^GCaMP/tdT^ mice. **(E)** Linear regression of the average z-score fluorescence ratio during food approach and consumption in relation to pellet events across the sessions of fasted LC^GCaMP/tdT^ mice. Approach (Slope=-0.02126+0.006555; *R*^*2*^=0.02569; *F*_1,399_=10.52, \*\**P*<0.01) and Consumption (Slope=0.01562+0.004806; *R*^2^=0.02579; *F*_1,399_=10.56, \*\**P*<0.01). **(F)** Average velocity during feeding-related behaviors. Two-way repeated measures ANOVA, main effect of behavior: *F*_3,36_=26.9, *P*<0.001, nutritional state x behavior interaction: *F*_3,36_=4.457, *P*=0.0092. Bonferroni post-hoc test, *P*<0.001 consumption (group b) vs. all other behavioral states (group a). Data are mean ± SEM. *n*=13 LC^GCaMP/tdT^ mice.

### Visual-Evoked LC-NE Activity is Attenuated in Sated Mice

Next, we wanted to determine if satiety state affects LC-NE responses to non-food stimuli. It is well established that LC neurons are activated by salient or unexpected sensory events*(12, 18)*, but whether this response is modulated by internal physiological states such as hunger is unknown. To address this question, we used fiber photometry to compare LC-NE activity evoked by a salient, 1-sec flash of light presented to mice that were fasted overnight or fed *ad libitum* **(Fig. 2A)**. In fasted LC^GCaMP/tdT^ mice, we observed a rapid, transient activation of LC-NE neurons to the visual stimulus **(Fig. 2B-C**). In mice fed *ad libitum*, however, the magnitude of LC-NE activity was smaller **(Fig. 2B-C)**, demonstrating that satiation attenuated LC-NE responses to visual stimuli. Importantly, movement velocity remained constant during visual stimulus presentation for fasted and sated mice **(Fig. 2D**), indicating that ambulation did not drive visual-evoked activity of LC neurons nor their response to satiety. To determine if LC response to visual stimuli habituated across the session, we next performed linear regression analyses for fasted and *ad libitum* fed mice presented with 30 light flashes across the 30-min session. Notably, we observed no change in light-evoked LC-NE activity across sessions for either the fasted or *ad libitum* fed mice **(Fig. 2E)**, indicating that the visual stimulus remained salient despite various presentations within the session. Collectively, our results demonstrate that internal satiety state affects LC-NE response to both food- and non-food related stimuli.

**Fig. 2.**
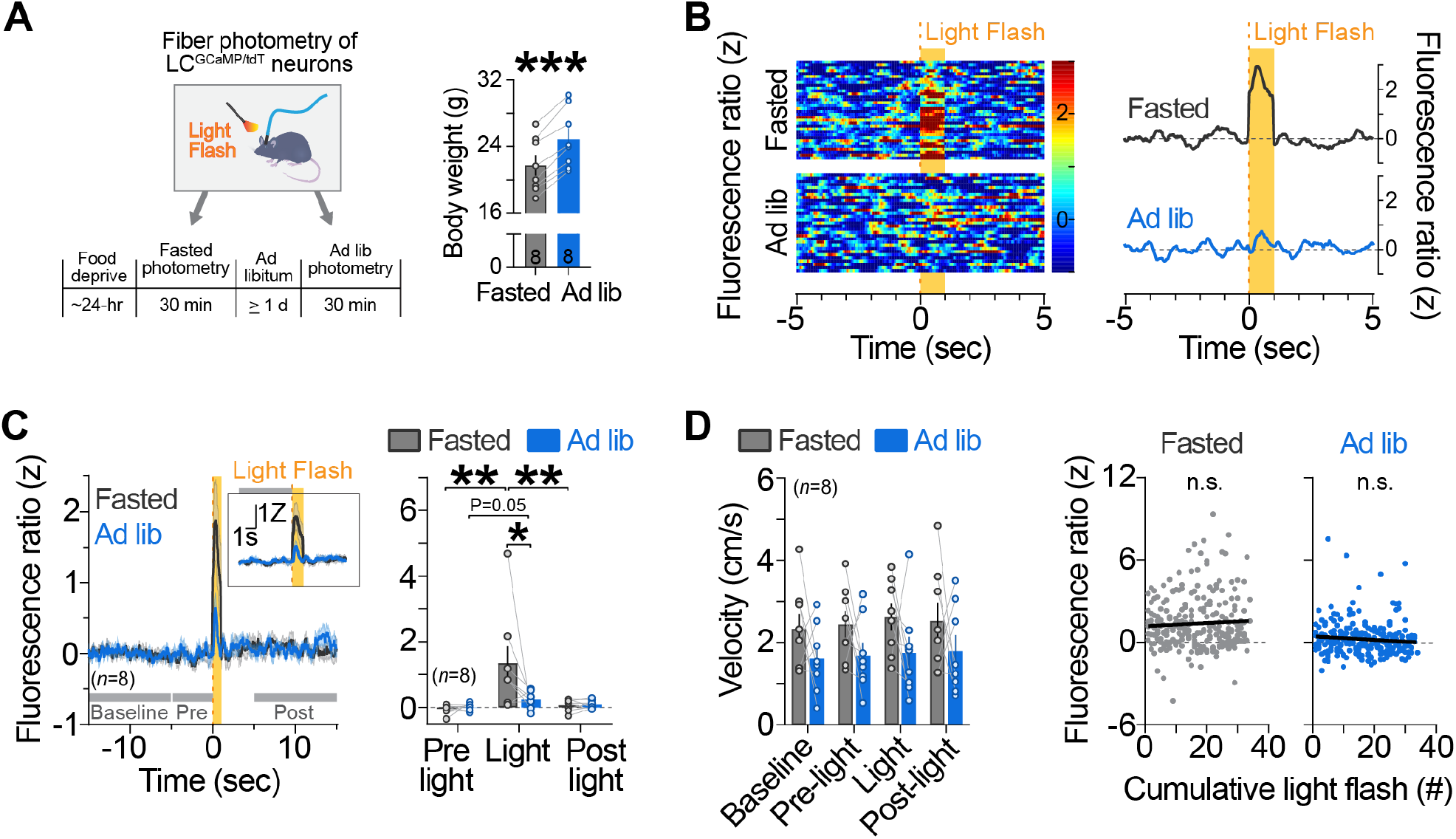
Visual-evoked LC-NE activity is influenced by nutritional state. **(A)** *Left*. Timeline of photometry recordings during presentation of light flashes. *Right*. Body weights during fasted and ad-libitum-fed recordings. Paired samples *t*-test, \*\*\**P*<0.001. Data are mean ± SEM. *n*=8 LC^GCaMP/tdT^ mice. **(B)** Fluorescence ratio expressed as a z-score aligned to visual stimulus during the fasted *(top)* and ad-libitum-fed *(bottom)* recordings from a representative LC^GCaMP/tdT^ mouse. Single trials are represented in the heatmap *(left)*, and average z-score fluorescence ratio is represented in the trace *(right)*. **(C)** Average z-score fluorescence ratio aligned to visual stimulus *(left)* and during visual-related events *(right)* in LC^GCaMP/tdT^ mice. Two-way repeated measures ANOVA, nutritional state x event interaction: *F*_2,14_=3.938, *P*=0.0440. Bonferroni post-hoc test, \*\**P*<0.01, \**P*<0.05. Paired samples *t*-test for pre-light vs. light in ad-libitum-fed mice: *t*_7_=2.33, *P*=0.0526. Data are mean ± SEM. *n*=8 LC^GCaMP/tdT^ mice. **(D)** Average velocity during visual-related events. Two-way repeated measures ANOVA, main effect of nutritional state: *F*_1,7_=1.728, *P*=0.2301, main effect of event: *F*_3,21_=2.235, *P*=0.1140, nutritional state x event interaction: *F*_3,21_=0.1346, *P*=0.9383. Data are mean ± SEM. *n*=8 LC^GCaMP/tdT^ mice. **(E)** Linear regression of the average z-score fluorescence ratio during light flashes across the fasted and ad-libitum-fed sessions of LC^GCaMP/tdT^ mice. Fasted (Slope=0.01118+0.01273; *R*^*2*^=0.002992, *F*_1,257_=0.7714, *P*=0.3806) and ad-libitum-fed (Slope=-0.01268+0.008154; *R*^2^=0.009470; *F*_1,253_=2.419, *P*=0.1211).

### Brief Stimulation of LC-NE Neurons is Sufficient to Suppressed Feeding in Fasted Mice

Given our finding that endogenous activity of LC-NE neurons is attenuated during feeding, we next sought to determine if feeding would be suppressed by brief stimulation of LC-NE neurons during the 10-sec of feeding immediately following pellet retrieval. To test this hypothesis, we injected Flp-dependent AAVs expressing the red-shifted channelrhodopsin variant ChrimsonR-tdT*(19, 20)* or tdT control in the LC of *Dbh*^*Flpo*^ mice*(16)* **(Fig. 3A)**. LC^ChrimsonR^ and LC^tdT^ mice were trained to eat grain pellets from the FED, fasted overnight and then behaviorally monitored while brief photostimulation was paired in real-time with feeding **(Fig. 3B)**. We found that LC^ChrimsonR^ mice consumed less during brief photostimulation compared to controls (**Fig. 3C)**. There was no difference between LC^ChrimsonR^ and LC^tdT^ mice in the latency to begin feeding or the number of pellets retrieved and dropped during brief photostimulation (**Fig. 3D, Fig. S1)**. These findings demonstrate that brief, behaviorally locked activation of LC-NE neurons is sufficient to suppress feeding in hungry mice.

**Fig. 3.**
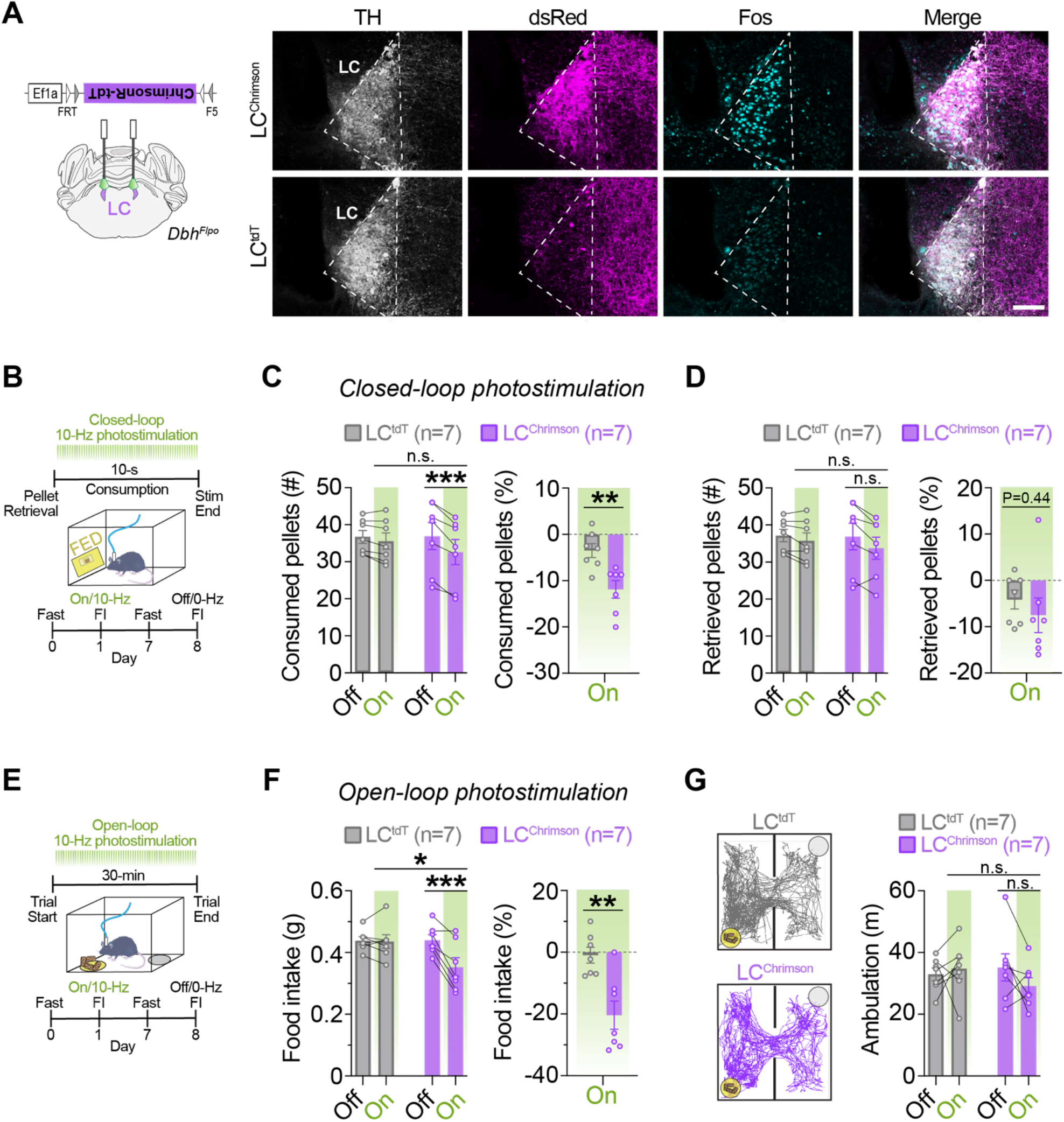
Optogenetic stimulation of LC-NE cell bodies suppresses feeding. **(A)** *Left*. Schematic illustration of coronal mouse brain showing location of Flp-dependent AAV used to drive ChrimsonR-tdT expression. *Right*. Immunofluorescent labeling of Fos (turquoise) in noradrenergic (TH, light gray), tdT-expressing (dsRed, magenta) locus coeruleus neurons in coronal brain sections from a representative LC^tdT^ and LC^ChrimsonR^ mouse following photostimulation (560-nm, 10-Hz, 10-ms) for 30-min. Scale, 100-μm. **(B)** Timeline of closed-loop optogenetic experiment wherein photostimulation (10-Hz, 10-ms pulses) was triggered upon pellet retrieval and occurred briefly for 10-sec during consumption from the feeding experimentation device (FED). **(C-D)** *Left*. Average feeding behavior in fasted mice during the closed-loop optogenetic experiment. Two-way repeated measures ANOVA, stimulation x virus interaction: number of pellets consumed (**C**, *F*_1,12_=13.70, *P*=0.0030) and number of pellets retrieved (**D**, *F*_1,12_=1.571, *P*=0.2339). Bonferroni post-hoc test, \*\*\**P*<0.001, \*\**P*<0.01. n.s., non-significant. Data are mean ± SEM. *n*=7 LC^tdT^ mice, *n*=7 LC^ChrimsonR^ mice. *Right*. Feeding behavior during photostimulation (On) as percent change from no stimulation (Off) during closed-loop optogenetic experiment in fasted mice. Unpaired samples *t*-test: percent change in pellets consumed (**c**, *t*_12_=3.504, \*\**P*<0.01) and percent change in pellets retrieved (**D**, *t*_12_=0.7904, *P*=0.4446). Data are mean ± SEM. *n*=7 LC^tdT^ mice, *n*=7 LC^ChrimsonR^ mice. **(E)** Timeline of food intake (FI) during open-loop photostimulation (10-Hz, 10-ms pulses). **(F)** *Left*. Average 30-min food intake measured in the presence (On) or absence (Off) of open-loop photostimulation in fasted mice. Two-way repeated measures ANOVA, stimulation x virus interaction: *F*_1,12_=14.42, *P*=0.0025. Bonferroni post-hoc test, \*\*\**P*<0.001, \*\**P*<0.01, \**P*<0.05. *Right*. Food intake during photostimulation as a percent change from no photostimulation. Unpaired samples *t*-test, *t*_12_= 3.716, \*\**P*<0.01. Data are mean ± SEM. *n*=7 LC^tdT^ mice, *n*=7 LC^ChrimsonR^ mice. **(G)** Ambulation in the FI task during open-loop photostimulation. *Left*. Representative traces show ambulation, yellow circle indicates the food cup location, and gray circle indicates the location of an empty cup. *Right*. Average ambulation in the FI task in fasted mice. Two-way repeated measures ANOVA, stimulation x virus interaction: *F*_1,12_=2.149, *P*=0.1684. n.s., non-significant. Data are mean ± SEM. *n*=7 LC^tdT^ mice, *n*=7 LC_ChrimsonR_ mice.

### Stimulation of LC-NE Rapidly Suppressed Feeding in Fasted Mice

Next, we sought to determine the effects of optogenetic stimulation of LC-NE neurons over a longer period on feeding. LC^ChrimsonR^ and LC^tdT^ control mice were overnight fasted, and intake of standard chow was measured for 30-min in the presence or absence of 560-nm optical pulses (10-Hz, 10-ms) while in a familiar arena **(Fig. 3E)**. We found LC^ChrimsonR^ mice had reduced food intake during optical stimulation compared to controls **(Fig. 3F)**, demonstrating that activating LC-NE neurons suppresses feeding. Importantly, there was no significant difference in ambulation between LC^ChrimsonR^ and LC^tdT^ controls **(Fig. 3G**), indicating LC-mediated suppression of feeding can occur independently from changes in ambulation. Taken together, our findings demonstrate that feeding can be suppressed by brief, behaviorally locked activation of LC-NE neurons as well as longer durations of optogenetic stimulation.

### Activation of LC-NE Suppressed Feeding Without Affecting Metabolism

To determine the metabolic effects of activating LC-NE neurons, we used triple transgenic *En1*^*cre*^; *Dbh*^*Flpo*^; *RC::FL-hM3Dq* (LC^hM3Dq^) mice expressing the excitatory Gq-coupled receptor hM3Dq fused to mCherry *(21)* **(Fig. 4A)**. This previously published non-viral intersectional approach allows selective and reproducible expression of hM3Dq in 99.6% of the anatomically defined LC located within the central gray*(22)*, and a small portion of the dorsal subcoeruleus and A7 immediately adjacent to and continuous with the LC*(16, 21, 23-25)*. To add to our prior characterization of the LC^hM3Dq^ mouse line we first performed *in vivo* fiber photometry in LC^hM3Dq^ mice expressing GCaMP6f and littermate control mice expressing GCaMP6f (LC^Controls/GCaMP6f^) or lacking GCaM6f expression (LC^Controls/EGFP^) following administration of clozapine N-oxide (CNO, 5 mg/kg i.p.) or vehicle. Recordings revealed a sustained increase in LC-NE activity following CNO treatment in LC^hM3Dq^ mice compared to controls **(Fig. S2)**. No effect of CNO or vehicle was observed in LC^Controls/GCaMP6f^ or LC^Controls/EGFP^ littermate control mice **(Fig. S2**). In all mice expressing GCaMP6f, but not GFP, we observed increases in LC activity during handling and injection **(Fig. S2**), validating the sensitivity of our recording conditions to detect an established threat-related response*(18, 26, 27)*.

**Fig. 4.**
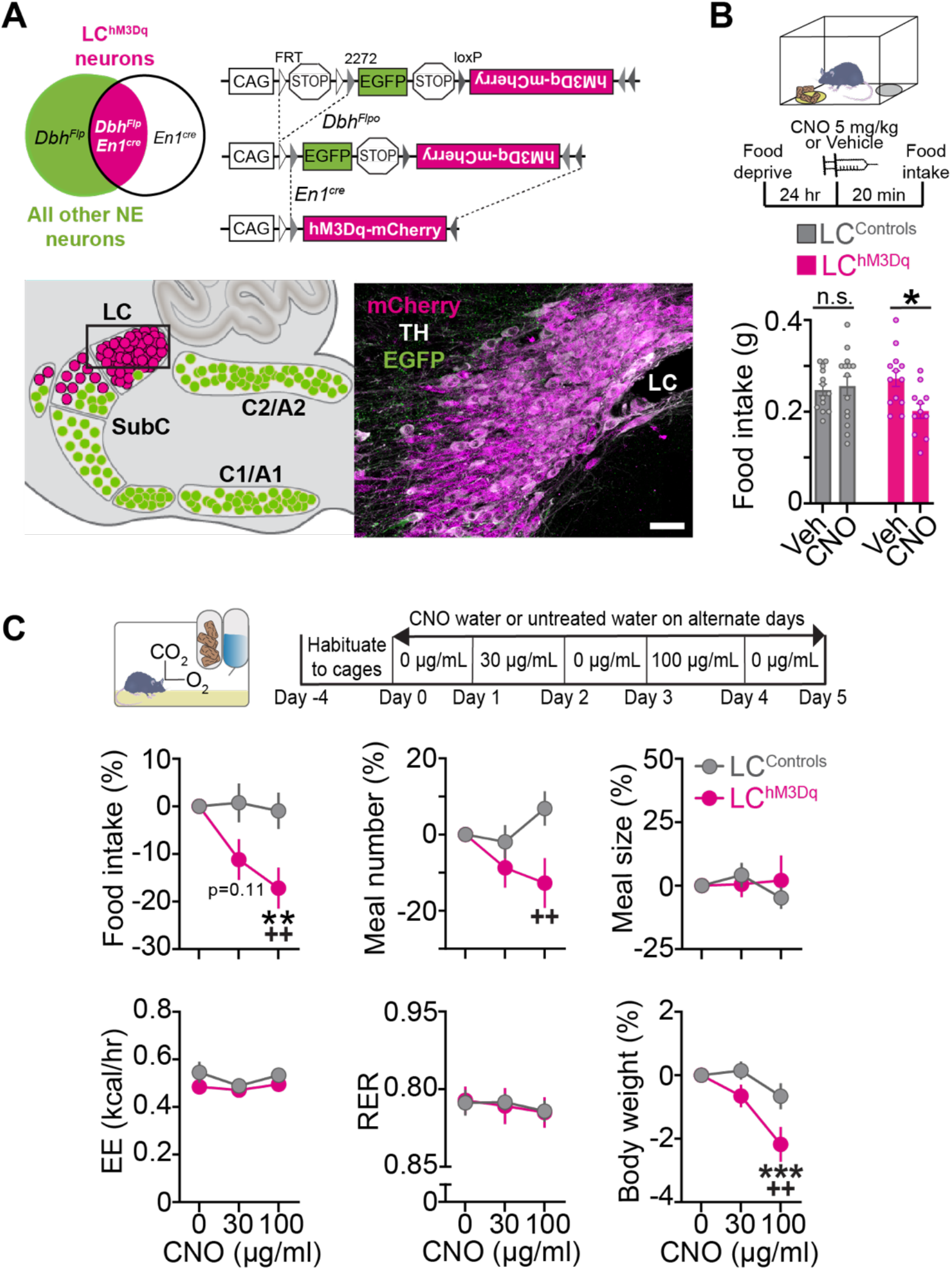
Chemogenetic activation of LC-NE neurons suppresses feeding without altering metabolism. **(A)** *Left*. Schematic illustration of intersectional genetic strategy. Recombination of *RC::FL-hM3Dq* allele by *Dbh*^*Flpo*^ and *En1*^*cre*^ results in hM3Dq-mCherry expression in LC-NE neurons. Recombination by *Dbh*^*Flpo*^ alone leads to EGFP expression. *Right*. Schematic of sagittal mouse hindbrain compressed across the mediolateral axis. Parasagittal section from LC^hM3Dq^ brain reveals hM3Dq-mCherry expression in LC-NE neurons. Scale, 50-µm. **(B)** *Top*. Timeline of food intake (FI) experiments in fasted mice. *Bottom*. Average FI in fasted mice. Two-way between-subjects ANOVA, drug x genotype interaction: *F*_1,47_=5.20, *P*=0.0272. Bonferroni post-hoc test, \**P*<0.05. n.s., non-significant. Data are mean ± SEM. *n*=13 vehicle- and *n*=14 CNO-treated LC^Controls^. *n*=13 vehicle and *n*=11 CNO-treated LC^hM3Dq^ mice. **(C)** *Top*. Timeline of CNO water at 30 and 100 µg/mL. *Bottom*. Behavioral and metabolic measures in the automated homecage. Two-way repeated measures ANOVA, drug x genotype interaction: food intake *(F*_2,54_=3.64, *P*=0.0329), meal number *(F*_2,54_=3.478, *P*=0.0379), meal size *(F*_2,54_=0.8542, *P*=0.4313), energy expenditure (EE, *F*_2,54_=0.4781, *P*=0.6226), respiratory exchange rate (RER, *F*_2,54_=0.04714, *P*=0.9540), and body weight *(F*_2,70_=3.647, *P*=0.0312). Bonferroni post-hoc test, \*\*\**P*<0.001, \*\**P*<0.01 vs. vehicle; ^++^*P*<0.01 vs. LC^Controls^. Data are mean ± SEM. *n*=20 LC^Controls^, *n*=9 LC^hM3Dq^ mice for all measures except body weight wherein *n*=17 LC^hM3Dq^ mice.

To confirm that our chemogenetic strategy for activation of LC-NE neurons would suppress feeding in hungry mice, LC^hM3Dq^ mice and littermate controls were overnight fasted and given an i.p. dose of CNO or vehicle before feeding was measured in a novel arena **(Fig. 4B)**. We found CNO suppressed feeding in LC^hM3Dq^ mice but not littermate controls **(Fig. 4B)**, demonstrating that chemogenetic activation of LC-NE neurons suppresses feeding in hungry mice. To determine precisely how LC activation influences daily feeding and metabolism, LC^hM3Dq^ and littermate control mice were housed for several days in the Labmaster homecage, which tracks meals and metabolic parameters by indirect calorimetry (energy expenditure and respiratory exchange rate) **(Fig. 4C, Fig. S3)**. To avoid the stress induced by daily injection of CNO, plain water or CNO (30 or 100 µg/mL) in the drinking water was administered on alternate days. In CNO-treated LC^hM3Dq^ mice, activation of LC-NE neurons dose-dependently suppressed feeding by reducing the number of meals consumed, without affecting meal size **(Fig. 4C, Fig. S3)**. These changes resulted in weight loss in CNO-treated LC^hM3Dq^ mice **(Fig. 4C)**. Analysis of circadian behavior revealed that suppression of feeding and drinking occurred specifically during lights-off (when LC^hM3Dq^ mice drank the most CNO), and this effect was reversible during lights-on (when mice drank less CNO) and upon removal of the drug the next day **(Fig. S4A-B)**, indicating that LC-mediated suppression of ingestive behavior was specific and reversible. Further, using indirect calorimetry, we found CNO had no effect on energy expenditure or respiratory exchange rate in LC^hM3Dq^ mice and littermate controls **(Fig. 4C)**. Importantly, the average daily dose of CNO was similar for LC^hM3Dq^ mice and littermate controls (**Fig. S4C**). These findings collectively demonstrate that activation of LC-NE neurons suppresses feeding without altering metabolism, and therefore we focused our efforts to measures of food intake in subsequent experiments.

### Inhibition of LC-NE Neurons has No Significant Effect on Acute Food Intake

Given that LC-NE activation suppresses feeding, we next sought to determine if inhibition of LC-NE neurons would promote feeding. To test this hypothesis, we injected a cre-dependent AAV expressing the Gi-coupled receptor hM4Di-mCherry*(28)* or mCherry in the LC of *Dbh*^*cre*^ mice^*(29)*^**(Fig. S5A)**. To confirm the expressed hM4Di receptor was functional, we replicated a prior chemogenetic study*(30)* and found that inhibiting LC-NE neurons attenuates restraint stress-induced anxiety-like behavior in the open field, without affecting total ambulation (**Fig. S5B)**. To determine if LC-NE activity is required for feeding, LC^hM4Di^ mice and controls were overnight fasted and treated with CNO (1 mg/kg i.p.) or vehicle before placement in a familiar arena containing standard chow **(Fig. S5C)**. No significant change in feeding was observed between CNO-treated LC^hM4Di^ mice and LC^mCherry^ controls (**Fig. S5C)**, suggesting that LC-NE activity is not necessary for promoting feeding in hungry mice.

### Stimulation of the LC-Lateral Hypothalamus Circuit Suppresses Feeding, Elicits Aversion, and Enhances Anxiety-Like Behavior

Prior studies have shown that LC-NE neurons project to the lateral hypothalamus area (LHA)*(16, 30, 31)* and that NE has a strong inhibitory effect when applied directly in the LHA*(32-36)*. To determine if the LC may mediate this effect, we measured Fos immunoreactivity in LC^hM3Dq^ mice following treatment of CNO (1 mg/kg i.p.) or vehicle. Activation of LC-NE neurons resulted in a significant reduction in Fos expression in the LHA **(Fig. S6A)**. To determine if this response was specific to the LHA, we measured Fos expression in two additional feeding-related targets of the LC, the dorsal medial and ventromedial hypothalamic nuclei. We observed no change in Fos expression in either nucleus **(Fig. S6B-C)**.

To directly test whether stimulation of the LC-LHA circuit suppresses feeding, we injected AAVs expressing cre-dependent channelrhodopsin-2 (ChR2)*(37, 38)* or EYFP in the LC of *Dbh*^*cre*^ mice*(29)* **(Fig. 5A)**. Consistent with prior observations*(16, 30, 31)*, we observed LC derived axons in the LHA **(Fig. 5B-C)**. We next measured food intake of overnight fasted LC-LHA^ChR2^ and control mice in the presence or absence of 465-nm optical pulses (10-Hz, 10-ms) for 30-min while in a familiar arena (**Fig. 5D)**. Optical stimulation suppressed feeding in LC-LHA^ChR2^ mice without affecting ambulation (**Fig. 5E-F)**. Importantly, no change in these behaviors was observed in LC-LHA^EYFP^ controls following optical stimulation (**Fig. 5E-F)**. Our findings demonstrate that activation of the LC-LHA noradrenergic pathway is sufficient to suppress feeding in hungry mice.

**Fig. 5.**
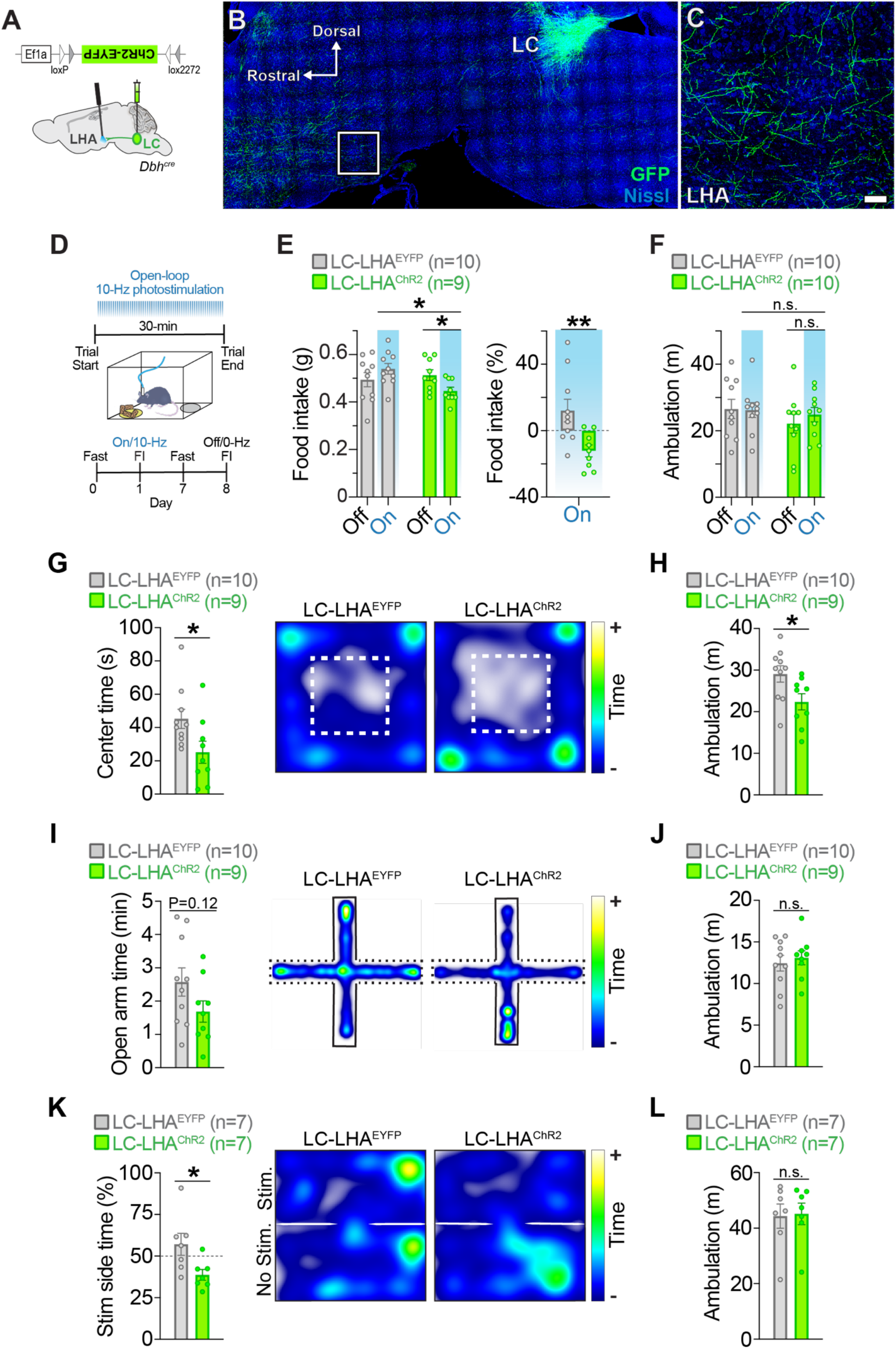
Optogenetic stimulation of the LC-LHA noradrenergic pathway suppresses feeding, elicits aversion, and enhances anxiety-like behavior. **(A)** Schematic illustration of sagittal mouse brain shows location of cre-dependent AAV used to drive ChR2-EYFP expression and location of fiberoptic probes. **(B-C)** Parasagittal brain section from a *Dbh*^*cre*^; LC-LHA^EYFP^ mouse shows restricted EYFP expression in LC-NE neurons. High magnification image shows EYFP-expressing LC-NE axonal projections in the LHA. Scale is 400-µm (brain) and 150-µm (LHA). **(D)** Timeline of food intake (FI) during open-loop photostimulation (465-nm, 10-Hz, 10-ms pulses). **(E-F)** Average feeding-related behaviors measured in the presence (On) or absence (Off) of open-loop photostimulation in fasted mice. Two-way repeated measures ANOVA, stimulation x virus interaction: food intake (**e** *Left, F*_1,17_=10.8, *P*=0.0044) and ambulation (**F**, *F*_1,18_=0.655, *P*=0.4290). Bonferroni post-hoc test, \**P*<0.05. n.s., non-significant. Data are mean ± SEM. *n*=10 LC-LHA^EYFP^ mice, *n*=9-10 LC-LHA^ChR2^ mice. Unpaired samples *t*-test: food intake during photostimulation as a percent change from no photostimulation (**e** *Right, t*_17_= 3.055, \*\**P*<0.01). Data are mean ± SEM. *n*=10 LC-LHA^EYFP^ mice, *n*=9 LC-LHA^ChR2^ mice. **(G-H)** OFT behaviors. Unpaired samples *t*-test: center time (**G**, *t*_17_=2.279, \**P*<0.05) and ambulation (**H**, *t*_17_=2.411, \**P*<0.05). Data are mean ± SEM. *n*=10 LC-LHA^EYFP^ mice, *n*=9 LC-LHA^ChR2^ mice. **(I-J)** EPM behaviors. Unpaired samples *t*-test: open arm time (**I**, *t*_17_=1.638, *P*=0.1199) and ambulation (**J**, *t*_17_=0.5101, *P*=0.6165). Data are mean ± SEM. *n*=10 LC-LHA^EYFP^ mice, *n*=9 LC-LHA^ChR2^ mice. **(K-L)** RTPT behaviors. Unpaired samples *t*-test: stimulation side time (**K**, *t*_12_=2.498, \**P*<0.05) and ambulation (**L**, *t*_12_=0.1369, *P*=0.8934). Data are mean ± SEM. *n*=7 LC-LHA^EYFP^ mice, *n*=7 LC-LHA^ChR2^ mice. **(G, I, K)** Representative spatial location heatmaps show time spent exploring the arenas.

We next sought to determine if stimulating the LC-LHA pathway would elicit negative affective behaviors associated with stress, as it is well-known that stressful events activate LC-NE neurons*(39, 40)* and stimulating these neurons mimics stress by producing anxiety and aversion*(21, 30, 41)*. To measure anxiety-like behavior, we ran LC-LHA^ChR2^ and LC-LHA^EYFP^ mice in the open field test (OFT) and elevated plus maze (EPM) during optical stimulation. We observed LC-LHA^ChR2^ mice spent significantly less time in the center of the OFT **(Fig. 5G)** and tended to spend less time in the open arms of the EPM **(Fig. 5I)**, demonstrating that activation of the LC-LHA noradrenergic pathway is anxiogenic. To assess if stimulation of the LC-LHA pathway has a negative or positive valence, we employed a real-time place preference test (RTPT) that triggers photostimulation upon entry into a designated side of the arena. We found LC-LHA^ChR2^ mice spent less time in the stimulation-paired side compared to LC-LHA^EYFP^ controls **(Fig. 5K)**, indicating an aversive behavioral response resulting from LC-LHA circuit activation. During photostimulation, LC-LHA^ChR2^ mice had reduced ambulation in the OFT compared to controls, but no change in the EPM and RTPT **(Fig. 5H, J, L)**, suggesting that the LC-LHA circuit does not have an overall impact on locomotion. Subsequent assessment of Fos-immunoreactivity revealed that photostimulation of ChR2-expressing LHA terminals did not induce antidromic activity of LC-NE neurons **(Fig. S7)**. These findings demonstrate that activating the LC-LHA pathway elicits negative affect behaviors and suppresses feeding when photostimulation is pulsed throughout the behavioral paradigm.

### Stimulation of the LC-LHA Circuit Suppresses Feeding When Stimulation is Pulsed for Longer Durations, Not When Briefly Paired with Feeding

To test if feeding would be suppressed by selective activation of the LC-LHA pathway during consumption, we injected AAVs expressing Flp-dependent ChrimsonR-tdT*(19, 20)* or tdT in the LC of *Dbh*^*Flpo*^ mice **(Fig. 6A)**. LC-LHA^ChrimsonR^ mice and controls were trained to eat grain-based pellets from the FED, overnight fasted, and feeding was measured while 560-nm optical pulses (10-Hz, 10-ms) were delivered for 10-sec during consumption **(Fig. 6B)**, when LC-NE neurons are endogenously less active **(Fig.1C-D)**. We observed no significant change in the number of pellets consumed or retrieved between LC-LHA^ChrimsonR^ mice and controls (**Fig. 6C-D)**, suggesting that feeding is unaffected by brief, behaviorally locked activation of the LC-LHA pathway. In a separate experiment delivering 30-min of stimulation (10-Hz, 10-ms), we found this longer duration of pulsing suppressed food intake and ambulation in LC-LHA^ChrimsonR^ mice (**Fig. 6E-G)**, reproducing the feeding effect using LC-LHA^ChR2^ mice. Together, our findings demonstrate that stimulating the LC-LHA pathway for longer durations suppresses feeding potentially due to an enhancement of negative valence and reduction in locomotor activity.

**Fig. 6.**
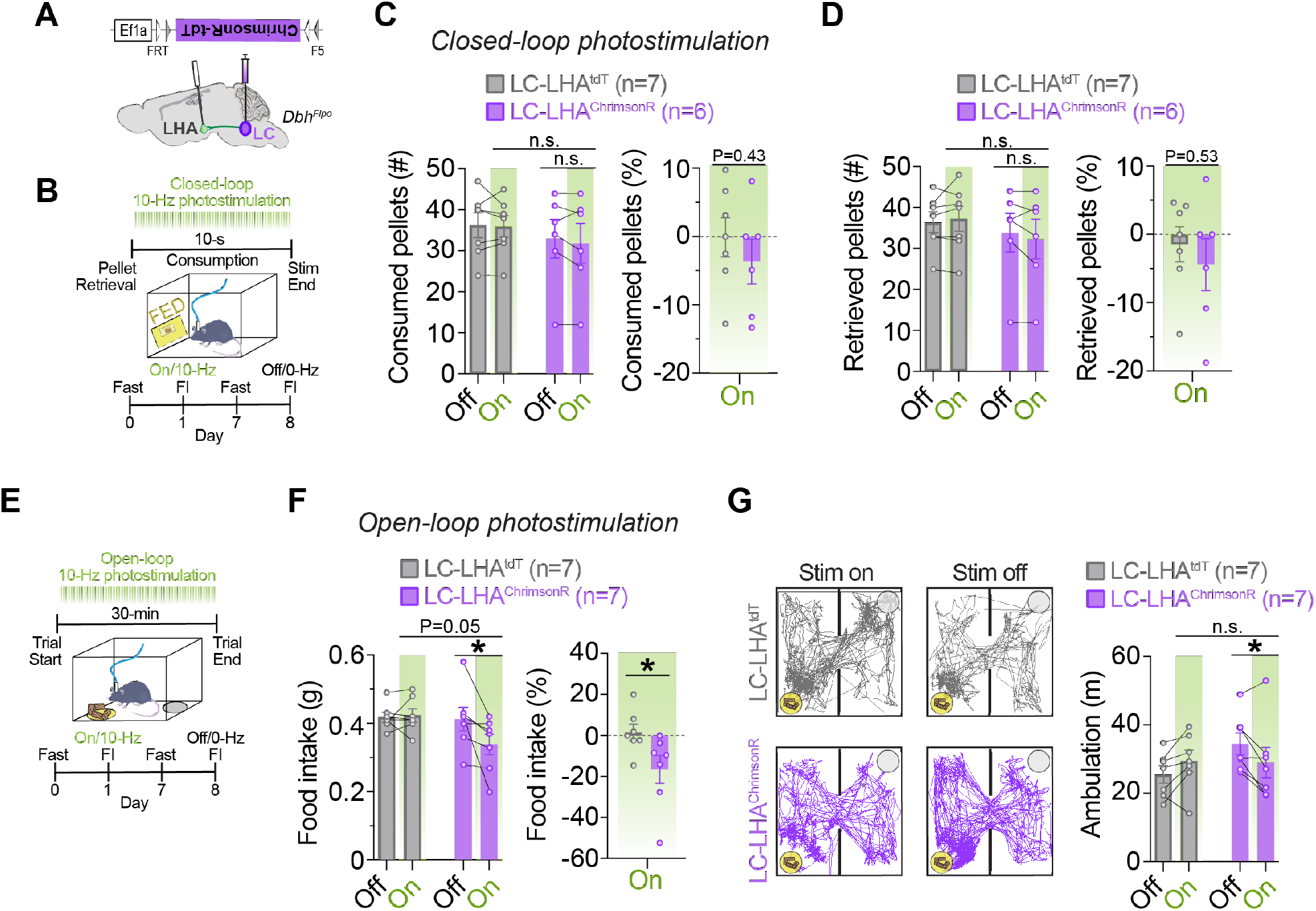
Optogenetic stimulation of the LC-LHA noradrenergic pathway suppresses feeding when stimulation is pulsed during the entire behavioral assay, but not when restricted to feeding events. **(A)** Schematic illustration of coronal mouse brain shows location of Flp-dependent AAV used to drive ChrimsonR-tdT expression and location of fiberoptic probes. **(B)** Timeline of closed-loop optogenetic experiment wherein photostimulation (560-nm, 10-Hz, 10-ms pulses) was triggered upon pellet retrieval and occurred briefly for 10-sec during consumption from the feeding experimentation device (FED). **(C-D)** *Left*. Average feeding behavior during the closed-loop optogenetic experiment in fasted mice. Two-way repeated measures ANOVA, stimulation x virus interaction: number of pellets consumed (**c**, *F*_1,11_=0.2807, *P*=0.6068) and number of pellets retrieved (**d**, *F*_1,11_=1.857, *P*=0.2003). n.s., non-significant. Data are mean ± SEM. *n*=7 LC-LHA^tdT^ mice, *n*=6 LC-LHA^ChrimsonR^ mice. *Right*. Feeding behavior during photostimulation (On) as percent change from no stimulation (Off) during the closed-loop optogenetic experiment in fasted mice. Unpaired samples *t*-test: percent change in pellets consumed (**c**, *t*_11_=0.8160, *P*=0.4318) and percent change in pellets retrieved (**D**, *t*_11_=0.6549, *P*=0.5260). Data are mean ± SEM. *n*=7 LC-LHA^tdT^ mice, *n*=6 LC-LHA^ChrimsonR^ mice. **(E)** Timeline of food intake (FI) during open-loop photostimulation (10-Hz, 10-ms pulses). **(F)** *Left*. Average 30-min food intake measured in the presence (On) or absence (Off) of open-loop photostimulation in fasted mice. Two-way repeated measures ANOVA, stimulation x virus interaction: *F*_1,12_=5.116, *P*=0.0431. Bonferroni post-hoc test, \**P*<0.05. Data are mean ± SEM. *n*=7 LC-LHA^tdT^ mice, *n*=7 LC-LHA^ChrimsonR^ mice. *Right*. Food intake during photostimulation (On) as a percent change from no photostimulation (Off). Unpaired samples *t*-test, *t*_12_=2.269, \**P*<0.05. Data are mean ± SEM. *n*=7 LC-LHA^tdT^ mice, *n*=7 LC-LHA^ChrimsonR^ mice. **(G)** Ambulation in the FI task during open-loop photosimulation. *Left*. Representative traces show ambulation, yellow circle indicates the food cup location, and gray circle indicates the location of an empty cup. *Right*. Average 30-min ambulation in the FI assay during open-loop photostimulation in fasted mice. Two-way repeated measures ANOVA, stimulation x virus interaction: *F*_1,12_=10.33, *P*=0.0074. Bonferroni post-hoc test, \**P*<0.05. n.s., non-significant. Data are mean ± SEM. *n*=7 LC-LHA^tdT^ mice, *n*=7 LC-LHA^ChrimsonR^ mice.

### Inhibition of the LC-LHA Pathway has No Significant Effect on Acute Food Intake

To test if inhibition of the LC-LHA circuit enhances feeding, *Dbh*^*cre*^ mice^*(29)*^ were injected with AAVs expressing a cre-dependent hM4Di*(28)* receptor or mCherry and implanted with bilateral cannula above the LHA **(Fig. S5D**). We overnight fasted LC-LHA^hM4Di^ mice and controls, delivered CNO (0.3 µM) or vehicle directly into the LHA, and then measured food intake after 30-min. We observed no change in feeding between LC-LHA^hM4Di^ mice and controls administered CNO or vehicle into the LHA (**Fig. S5D)**. Further, we observed no change in feeding in sated mice using another strategy in which cre-dependent halorhodopsin (eNpHR3.0-EYFP) or EYFP was expressed in the LC of *Dbh*^*cre*^ mice^*(29)*^ **(Fig. S5E)**. Collectively, our opto- and chemogenetic studies demonstrate that activating the LC-LHA pathway suppresses feeding, but inhibiting the pathway does not promote feeding, indicating the LC-LHA pathway operates exclusively like a “brake” on feeding.

## DISCUSSION

It is well established that LC-NE neurons promote alerting and orienting to salient external stimuli*(1-6)*, but their involvement in the regulation of feeding remains less understood. In the current study we found that, in contrast to their response to sensory stimuli and stressors*(12, 18, 42, 43)*, endogenous activity of LC-NE neurons is suppressed during feeding. Interestingly, satiety attenuated the magnitude of LC-NE responses to both food and flash of light. Further, we found that chemogenetic activation of LC-NE neurons suppressed ingestive behavior without altering metabolism and yielded modest weight loss (2% loss). Further we found activation of LC-NE neurons reduced circadian food intake and body weight. Using optogenetics, we found that feeding is suppressed when LC-NE neurons are activated by either a brief stimulation paired with feeding or stimulation over a longer duration. Chemogenetic inhibition of LC-NE neurons did not promote feeding. This finding is consistent with prior results showing LC-NE activation promotes stress-like anxiety responses, but inhibition has no effect on baseline anxiety*(30)*. Lastly, we identified a LC-LHA circuit that is sufficient but not necessary for the suppression of feeding.

Our *in vivo* photometry experiments demonstrate that the activity pattern of LC-NE neurons is dynamically modulated during the consummatory sequence, with initial activation during food approach followed by a suppression of LC activity during consumption. This native pattern of LC-NE activity to food is unlikely attributed to changes in sleep, learning, stress, or anxiety, as our experimental mice were habituated to all aspects of the assay and were tested during the portion of the circadian cycle when they are the most active. Instead, the approach-related LC response we observed likely reflects an appetitive response to food in hungry mice. This finding is in agreement with prior electrophysiological studies that show burst firing of LC neurons is associated with Pavlovian appetitive behavior in monkeys (e.g., lipping)*(44)*. Given LC neurons are well-known to modulate arousal*(1, 2, 45, 46)*, the decrease in LC activity we observed during feeding may reflect a decrease in arousal and/or disengagement with external sensory inputs to facilitate consummatory behavior. Supporting this interpretation, our closed-loop optogenetic results show that activating LC-NE neurons during consumption attenuated food intake, demonstrating that transiently suppressing LC-NE activity is a key mechanism that is naturally involved in the regulation of feeding.

As satiation developed across the trial, there was a gradual decrease in spontaneous LC-NE activity during food approach, which may contribute to the decrease in salience of food as energy balance is restored. Additionally, the suppression of LC-NE activity during consumption was mildly attenuated when transitioning from hunger to satiety, which could redirect attention to the external environment as satiation develops, although additional research is needed to test this possibility. Further, the increase in visual-evoked LC-NE activity in fasted mice suggests that hunger potentiates LC responses to external sensory stimuli, which may potentially aid in food seeking and self-preservation during foraging. Such potential interpretation is congruent with integrative theories of LC-NE neurons which describe that this modulatory system is designed to optimize behavioral performance to a changing environment (as (reviewed in *1, 2, 4*, and *47)*). Our findings demonstrate a previously unrecognized role for nutritional state in modulating endogenous LC responses to food and sensory stimuli.

Recently it has been shown using *in vivo* calcium imaging that a subpopulation of glutamatergic neurons in the peri-LC (peri-LC^VGLUT2^) is inhibited during consumption in a manner attenuated by satiety state*(48)*, mimicking the activity patterns we observed in LC-NE neurons. It will be important for future studies to map the functional connectivity between LC-NE neurons and neighboring and distal targets that orchestrate behavioral states involved in feeding regulation. A recent study found that fear-induced suppression of feeding is mediated by LC-NE neurons through a projection to the lateral PBN*(13)*. In the current study, we found the LC→LHA pathway suppressed feeding while also enhancing aversion and anxiety-like behavior. We then went on to demonstrate that stimulating the LC-LHA pathway suppressed feeding when optical pulses were continuously delivered during the feeding assay (30-min), but not when transiently paired with consumption events (10-sec). This divergent effect may be related to the cellular, synaptic, and molecular mechanisms that remain to be determined, including the downstream synapses engaged and the type and amount of norepinephrine and/or neuropeptide(s) released from LC terminals following optical stimulation.

A series of seminal studies established that exogenous delivery of NE suppresses feeding*(49-52)*, and that this effect is mimicked by direct delivery of NE agonists into the LHA*(53-57*, see also *58)*. However, the source of NE that modulates these functions remains unclear, as the LHA receives input from LC and A2 noradrenergic nuclei*(16, 59-61)*. Using optogenetics, we have now revealed two previously unrecognized roles for a LC to LHA circuit: (1) the suppression of feeding and (2) enhancement of negative valence behavior (i.e., anxiety, aversion). This dual function is coherent with the well-established literature that shows LHA cell-types drive a variety of complex behaviors, including feeding, drinking, anxiety, and arousal*(62-66)*. Our results are also consistent with a recent finding that chemogenetic activation of the LC-LHA pathway suppresses binge-like ethanol intake*(67)*, suggesting this pathway may broadly regulate motivated behaviors. An important future direction will be to determine whether the feeding and anxiety-related changes following LC-LHA pathway stimulation are mediated by separate or overlapping signaling mechanisms. For example, distinct subsets of LC-NE neurons and/or LHA cell-types may underlie feeding and affective behavior. Given the widespread projections of LC-NE neurons throughout the brain*(16)*, projections beyond the hypothalamus are also likely involved in LC modulation of feeding.

Here we demonstrate that during feeding LC-NE activity is transiently suppressed, and that food intake can be attenuated by briefly activating LC-NE neurons. We also identified a previously unrecognized role for nutritional state in modulating endogenous LC responses to both food and visual flash of light. Further, our data provide emerging insight into the neural circuitry of motivated behavior, revealing a dual role of the LC-LHA pathway in feeding suppression and negative affective behavior. In context with the broader LC literature*(1, 2, 45, 46, 68)*, our findings suggest that LC-NE neurons are involved in the modulation of feeding by integrating both external cues (e.g., anxiogenic environmental cues) and internal drives (e.g., nutritional state).

## MATERIALS AND METHODS

### Animals

All procedures related to the use of animals were approved by the NIEHS Animal Care and Use Committee and were in accordance with the National Institutes of Health guidelines for the care and use of laboratory animals. Adult (>P60) male and female mice were used for all studies. *En1*^*cre*^, *Dbh*^*cre*^, *Dbh*^*Flpo*^, and *RC::FL-hM3Dq* mouse colonies^*(16, 21, 29, 69)*^ are maintained on a C57BL/6J background. Triple transgenic *En1*^*cre*^; *Dbh*^*Flpo*^; *RC::FL-hM3Dq* mice and littermate controls were generated by crossing *En1*^*cre*^ to double transgenic *Dbh*^*Flpo*^; *RC::FL-hM3Dq* mice. Mice were maintained on a reverse 12:12-hr light:dark cycle with lights off at 8 AM. All experiments occurred during the dark period of the circadian cycle. Mice had *ad libitum* access to water and standard food (NIH-31, Harlan, Madison WI), except when mice underwent overnight food-deprivation before the food intake test and photometry recordings where noted. Mice were group housed unless surgical implants or if CNO dosing by drinking water required single housing.

### Tissue Collection

Adult mice were deeply anesthetized with sodium pentobarbital and perfused transcardially with phosphate-buffered saline (PBS) followed by 4% paraformaldehyde in PBS (PFA/PBS). Brains were postfixed overnight by immersion in 4% PFA/PBS at 4°C. Following rinse in PBS, tissue was cryoprotected in 30% sucrose in PBS and embedded in Tissue Freezing Medium (General Data Healthcare). 40-μm free-floating coronal or sagittal brain sections were collected in PBS, transferred to a cryoprotectant, and stored at -80°C.

### Immunohistochemistry

For immunofluorescence staining, mCherry-expressing neurons were detected using Rat anti-mCherry primary antibody (1:1000) and Goat anti-rat Alexa Fluor 568 secondary antibody (1:1000). EGFP and EYFP-expressing neurons and axons were detected using Chicken anti-GFP primary antibody (1:10,000) and Goat anti-chicken Alexa Fluor 488 secondary antibody (1:1000). tdTomato+ neurons were detected using Rabbit anti-dsRed primary antibody (1:1000) and Goat anti-rabbit Alexa Fluor 568 secondary antibody (1:1000) or Rat anti-tdTomato primary antibody (1:2000) and Goat anti-rat Alexa Fluor 568 secondary antibody. The noradrenergic identity of neurons was confirmed with Rabbit anti-TH primary antibody (1:1000) and either Goat anti-rabbit 633 or Goat anti-rabbit 488 secondary antibody (1:1000). Mouse anti-TH primary antibody (1:500) was also used with Goat anti-mouse 488, Goat anti-mouse 568, or Goat anti-mouse 633 secondary antibody (1:1000). Fos was detected with Rabbit anti-cFos primary antibody (1:250, Santa Cruz, sc-52) and Goat anti-Rabbit 633 secondary antibody (1:1000). NeuroTrace 435/455 blue, fluorescent Nissl stain (1:50, N21479, ThermoFisher) was used to visualize neurons. For immunoperoxidase staining, Fos was detected using Rabbit anti-cFos primary antibody (1:2000, ab190289, Abcam) and biotinylated Goat anti-rabbit secondary antibody (1:500, BA-1000, Vector Labs) in conjunction with Vectastain Elite ABC kit and DAB substrate kit (Vector Labs). Coverslips were applied using Vectashield hard-set mounting medium with or without DAPI (H-1400 or H-1500, Vector Labs) or Prolong Diamond Anti-Fade mounting medium (P36970, Invitrogen). Methods were performed as previously described*(16, 21)*. Tissues from the different treatment groups were processed concurrently using the same batch of antibodies for all immunofluorescent experiments. The antibodies used are summarized in the ***Supplementary Materials***.

### Digital Image Processing

Images of immunofluorescently labeled sections were collected using Zeiss LSM780 or 880 inverted confocal microscopes (Carl Zeiss Inc., Oberkochen, Germany). When necessary, Zen Black 2012 Software (Carl Zeiss) was used to convert z-stacks to maximum intensity projections. Images were modified only by adjusting brightness and contrast across the entire image to optimize the fluorescence signal. Anatomical location was confirmed by reference to a mouse brain atlas*(70)*.

To measure Fos in LC-NE neurons, images of fluorescently stained sections were acquired at 20x using a Zeiss LSM780 and digital images were subsequently exported to MetaMorph (Molecular Devices, San Jose, CA). To measure Fos in the hypothalamus, brightfield images of DAB-stained brain sections were acquired at 40x using an Aperio AT2 slide scanner (Leica Biosystems Inc., Buffalo Grove, IL). Digital images were subsequently exported from Aperio Imagescope (Leica Biosystems) as an uncompressed *.tif file and opened in FIJI software*(71)* v2 for further analysis.

### Cell Counts

To measure Fos in LC-NE neurons, quantification was performed on every fourth 40-μm coronal section in the locus coeruleus and included at least 2-4 sections. An experimenter blind to treatment group performed the quantification. MetaMorph software was used to manually select TH+ and EGFP+ cells individually, and then we used the automated count nuclei feature to identify Fos cells using the following settings: approximate minimum width was 6-μm and maximum width was 9-μm with intensity above background of 10 gray levels. To determine colocalization, the AND operation was used within the Arithmetic process to count colocalized pixels that were ≥ than 20 μm^2^; this size was large enough to ignore background noise but small enough to count overlap of individual cells. To confirm our ability to detect Fos expression, we used LC^hM3Dq^ mice treated with CNO (5 mg/kg i.p.) and vehicle; this control tissue was run in every immunohistochemistry assay.

To measure Fos in hypothalamic nuclei, quantification was performed on every fourth 40-μm coronal section in the lateral, dorsomedial, and ventromedial hypothalamus, and included at least 2-4 sections. An experimenter blind to treatment group performed the quantification. FIJI software was used to measure the region-of-interest and perform automated counting. The number of Fos+ neurons was normalized to mm^2^ area.

### Drugs

For behavioral studies, clozapine n-oxide (CNO, 5-mg/kg or 1-mg/kg i.p., NIMH Drug Supply Program)*(21)* was administered 20-min prior to testing. For study of Fos expression, CNO (1-mg/kg i.p.) or vehicle was administered 2 hours before perfusion, and groups were treated identically (e.g., same experimenter for injection, same day of perfusion) and were not subjected to any other treatment (e.g., fasting) or behavioral assay prior to sacrifice. All compounds were injected at a volume of 0.1 mL/10 g body weight. Drugs were dissolved in DMSO (< 3%) and brought to volume using 0.9% physiological saline. For oral CNO administration, the doses selected were based on previously published papers (30 and 100 µg/mL)*(72-75)*. CNO water was prepared fresh daily by dissolving in 0.5% DMSO and brought to volume using reverse osmosis deionized (RODI) water. For *in vivo* microinfusion studies, bilateral intra-LHA microinjection of CNO (0.3-µM) was administered 30-min prior to testing at a dose previously shown to modulate feeding behavior upon local activation of DREADD-expressing neural circuits*(76)*. Microinjections of CNO were delivered in bubbled (95% O2 and 5% CO2) artificial cerebrospinal fluid (aCSF contained in mM: 182 sucrose, 20 NaCl, 0.5 KCl, 1 MgCl2-6H20, 1.2 NaH2PO4-H20, 26 NaHCO3, 10 glucose) at a volume of 100 nL at rate of 100 nL/min through an internal bilateral cannula.

### Viral Preparation

We generated a Flp-dependent GCaMP6f AAV construct for photometry experiments. pAAV-Ef1a-fDIO-EYFP (Addgene #55641) was digested with *Asc*I and *Nhe*I to remove the EYFP cDNA, and GCaMP6f cDNA was obtained from pGP-CMV-GCaMP6f (Addgene #40755)*(77)* digested with *Not*I and *Bgl*II restriction enzymes. After incompatible single-strand overhangs were filled-in to generate blunt ends, the two restriction fragments were ligated to produce plasmid pAAV-Ef1a-fDIO-GCaMP6f. To generate pAAV-Ef1a-fDIO-tdTomato, a tdTomato cDNA was isolated from pRSET-tdTomato (generously provided by Roger Tsien) and cloned into pAAV-Ef1a EYFP digested with *Asc*I and *Nhe*I. To generate pAAV-Ef1a-fDIO-ChrimsonR-tdTomato, a ChrimsonR-tdTomato fusion cDNA was isolated from pAAV-Syn-FLEX-rc[ChrimsonR-tdTomato] (Addgene #62723) and cloned into pAAV-Ef1a-fDIO EYFP digested as above. The viruses and titers that were used are summarized in the ***Supplementary Methods***.

### Surgery

Mice were anesthetized using 4% isoflurane and placed in a stereotaxic frame (Kopf Instruments, Model 900) equipped with a digital micromanipulator (Harvard Apparatus, Holliston, MA). Anesthesia was maintained with 0.5-2% isoflurane and/or a cocktail of ketamine/xylazine (100/7 mg/kg i.p.). Bupivicaine (270 µg in 0.1ml) was injected locally beneath the scalp prior to incision. For viral injections, 500-nL volume was delivered at the rate of 100-nL per min using a 30-gauge Neuros Hamilton syringe or 26-gauge needle attached to a microsyringe (7002, Hamilton, Reno NV) and pump (UMP3 UltraMicroPump, WPI, Sarasota FL). Needles were left in place for 5 minutes after infusion to minimize backflow of the virus upon withdrawal of the needle. To minimize post-operative pain, all mice received the analgesic Buprenorphine SR (1-mg/kg s.c.). Mice were given a minimum recovery time of 4 (for LC soma experiments) or 6 weeks (for LC fiber experiments) to allow sufficient time for viral infection, genetic recombination, and gene expression before behavioral measurements. See the ***Supplementary Materials*** for additional experimental details.

### LC-NE Photometry Recordings

*In vivo* optical recordings of GCaMP6f, tdTomato, and EGFP fluorescence intensities were measured in LC-NE neurons using a custom-built fiber photometry system, as previously described*(14, 15)*. Excitation light (488 nm, 20 mW continuous wave laser, OBIS 488LS-20, Coherent, Inc.) was launched into a fluorescence cube (DFM1, Thorlabs), reflected by a dichroic mirror (ZT488/561rpc-UF1, Chroma), and focused by an achromatic fiber port (PAFA-X-4-A, Thorlabs) onto the core of a multimode patch cable (M83L01 was used for 200-µm core fiber probes; M61L01 was used for 105-µm core fiber probes, Thorlabs). The distal end of the patch cable was connected to an optical fiber probe made with either a 200-µm core diameter, 0.39 NA multimode fiber (FT200EMT, Thorlabs) or a 105-µm core diameter, 0.22 NA multimode fiber (FG105LCA, Thorlabs) and a ceramic ferrule with a 1.25-mm OD (MM-CON2007–2300 or MM-CON2010-1270-2-WHT, Precision Fiber Products) by a ceramic sleeve (SM-CS125S, Precision Fiber Products Inc.). The power of the excitation light measured at the tip of the implantable fiber probe that was adjusted to 70 µW. Emitted fluorescence light was collected by the same optical fiber probe and patch cable, passed through the same dichroic mirror, and filtered through an emission filter (ZET 488/561m) before collected by a fiber port (PAF2S-11A, Thorlabs) and launched into a spectrometer (QE Pro-FL, Ocean Optics, Inc.) through a multi-mode patch cable (M200L02-A, 0.22 NA, AR-coated, 200/240 µm core/cladding, Thorlabs). Time-lapsed fluorescence emission spectra were visualized using Ocean View version 1.5. The spectrometer and camera (Basler, acA1300-60gm) were triggered using a Doric TTL pulse generator (OTPG_4, Neuroscience Studio Software, Doric Lenses, Inc., Quebec, Canada). Mice were given at least 1-week for surgical recovery and then were habituated for several days (20-min/day) to being tethered to optical patch cables while in the testing arena (30 × 30 × 35.5 cm, Phenotyper, Noldus Information Technology, Inc., Leesburg, VA, USA).

### Photometry Data Analysis

To quantify GCaMP6f emission and to separate fluorescence overlap between GCaMP6f and tdTomato, all raw emission spectra data were passed through a spectral linear unmixing algorithm written in R, as described previously*(14)*. To control for movement artifacts in the fluorescence signal (e.g., photon loss caused by tissue movement or bending of the fiber during mouse movement), the unmixed GCaMP6f coefficients were normalized to unmixed tdTomato coefficients to generate GCaMP6f/tdTomato fluorescence ratio values that were converted to z-scores. In a subset of mice without tdTomato expression (Fig. 4b), intensities of GCaMP6f or GFP emission were converted to fluorescence z-scores. See ***Supplementary Materials*** for additional details on the analysis.

### Behavioral Experiments

#### Chemogenetic Activation of LC-NE Neurons During Feeding

##### Food Intake Test (FI)

To motivate feeding, mice were fasted a day prior to the food intake test*(78)*. Food-deprived mice were allowed 20-min to explore a two-compartment arena (25 × 25 × 25 cm) that contained two petri-dishes (60 × 15 mm) located on opposite corners. One dish contained standard food (NIH-31) and the other dish was empty. The dish location was randomly assigned and counterbalanced for each treatment group. For drug experiments, mice received CNO (5-mg/kg i.p.)*(21)* or vehicle ∼20-min before the FI test. An experimenter blind to treatment recorded food intake. Mice were returned to *ad libitum* food after testing.

##### Labmaster Feeding and Metabolism

Mice were single-housed and allowed 4 days to acclimate to the Labmaster cages (23 × 13 × 1 cm) (TSE Systems, Bad Homburg, Germany). Following habituation, mice received water that contained vehicle or CNO (30 or 100 µg/mL) on alternate days, as used previously*(72-75)*. At the same circadian time each day (CT 10-11), food and water were replenished, and mice were weighed by an experimenter. Food spillage was also monitored and measured by an experimenter; wherein large spillage events (>1g in 15-min) were subtracted from total intake on rare occasions. *Ad libitum* food and drink was provided during the entire experiment. Measures of food and drink intake and gas exchange were collected every 15-min. Meal number and size was measured using the sequence meal analysis setting, whereby all meals were recorded chronologically to evaluate single feeding episodes. Meals started when food consumption was greater than 0.02 g and ended when consumption ceased for at least 5-min*(79, 80)*. For metabolic measures, O_2_ and CO_2_ were sampled for a 3-min period once every 15-min. Flow and sample rates were held at 0.3L/min and 0.25L/min respectively as determined by appropriate variation between the sample and reference cage. Respiratory exchange rate (RER) was derived by indirect calorimetry using the quotient VCO_2_/VO_2_. Energy expenditure (EE) was derived by the abbreviated Weir equation, EE=(3.941*VO_2_) + (1.106*VCO_2_). Metabolic values were normalized to the daily body weights (metabolic values/body weight). Daily cumulative intake and averages of gas exchange were assessed for each 24-hr drug-dosing day. Circadian data were summarized over three periods: early lights OFF (ZT11-20), lights ON (ZT20-8), and late lights OFF (ZT8-10). Data were collected using the Labmaster version 2.6.9.13409 (TSE Systems) and analyzed by an experimenter blind to treatment.

#### Optogenetic Stimulation of LC-NE Cell Bodies or LC-LHA Terminals During Feeding

LC^ChrimsonR^ mice and LC^tdT^ controls were briefly anesthetized (4% isoflurane) to connect optical implants targeting the bilateral LC or LHA to patch cables (0.37 NA cable measuring 0.5- or 0.6-meters in length, SBP(2)-200/220/900-0.37_0.45_FCM-2xMF1.25). Mice were given 20-min to recover in their homecage. During the experiment, optical stimulation (560-nm, 10-Hz, 10-ms pulse width, 6-7 mW total power from patch cable tip) was delivered using the Prizmatix Dual Channel Optogenetics-Lime-Green-LED. Behavior was video-recorded using a Basler (acA1300-60gm) or Logitech (C930e) camera and analyzed by an experimenter blind to treatment group.

##### Open-Loop Optogenetics During the Food Intake (FI) Test

All mice were habituated to being tethered in the testing arena (20 × 20 × 25 cm) for 20-min on a day prior to testing. Mice were tested in a 30-min session while receiving 560-nm optical stimulation (10-Hz, 10-ms pulse width) or no stimulation. Food intake was recorded by an experimenter blind to treatment.

##### Closed-Loop Optogenetics During the FED Pellet Intake Test

Prior to recordings, mice were habituated for several days to eating from the feeding experimentation device (FED)*(17)* in both the homecage and testing arena (Phenotyper, Noldus). Mice were then tested in a 60-min session while they received pulses (10-ms) of optical stimulation (560-nm, 10-Hz) that were triggered upon pellet retrieval and terminated 10-sec later. Thus, this protocol selectively delivered optical stimulation during discrete feeding events that were spontaneously initiated by the mice. Pellet retrieval timestamps were obtained from EthoVision, and retrieval and consumption events were confirmed by visual inspection of the video files. For quantification of other feeding behaviors, an experimenter blind to treatment condition used the manual score feature in EthoVision v15 to mark each experimental video file for precise timeframes in which the (1) mouse began consuming a pellet and (2) mouse dropped a pellet. “Consumption start” was defined as the first video frame in which a mouse brought a pellet up to its mouth. Consumption events that were difficult to see due to the positioning of the mouse or where the pellet was dropped prior to consumption were not scored for this measure (<10% of total consumption events for all mice). “Latency to consume” was calculated by subtracting consumption start time from pellet retrieval time. “Dropped pellets” was defined as any pellet that was dropped during the experiment, regardless of whether that pellet was eaten after dropping.

#### Optogenetic Stimulation of LC-LHA Circuit During Feeding & Anxiety-Related Behaviors

LC-LHA^ChR2^ mice and LC-LHA^EYFP^ controls were briefly anesthetized (4% isoflurane) to connect the fiber-optic cannula to bilateral optical cables (0.48 NA cable measuring 0.5-, 0.6- or 1-meter in length with LC ferrule, BFP(2)_200/300/900-0.48_FCM-2xMF1.25). While in the homecage, mice were given 20-min to recover before optical stimulation. For stimulation of LC fibers, mice received 10-Hz (10-ms pulse width) photostimulation (6-9 mW total power) using the Doric system (LEDFRJ 465 nm powered by LEDD driver, Neuroscience Studio Software, Doric Lenses), as previously described*(30, 81)*. Behavior was video-recorded using a Basler (acA1300-60gm) or Logitech (C930e) camera and analyzed by an experimenter blind to treatment group.

##### Food Intake Test (FI)

All mice were habituated to being tethered in the testing arena (20 × 20 × 25 cm) for 20-min on a day prior to testing. Mice were tested in a 30-min session while receiving photostimulation (10-Hz, 10-ms pulse width). Food intake was recorded by an experimenter blind to treatment.

##### Elevated Plus Maze (EPM)

Mice were placed in a ‘+’ shaped maze (Stoelting Co), as previously described*(21)*. The maze had two open (35 × 5 cm, 15 mm lip) and two closed arms (35 × 5 × 15 cm) and was elevated 50-cm. Mice could explore for 5-min while receiving photostimulation (10-Hz, 10-ms pulse width). The time spent in the open arms (minutes) and ambulation (meters) were recorded using EthoVision v12-14. Open arm time is reported as (open arm time) / (open arm time + closed arm time) x 100).

##### Real-Time Place Test (RTPT)

Mice were allowed 20-min to explore an unbiased two-compartment arena (50 × 50 × 25 cm), as described previously*(30, 78)*. Photostimulation (10-Hz, 10-ms pulse width) occurred upon entry into one compartment and persisted until the mouse exited the compartment. The stimulation compartment was randomly assigned and counterbalanced for each treatment group. The percentage of time spent in the photostimulation-paired compartment and ambulation were recorded by EthoVision v12-14.

##### Open Field Test (OFT)

Mice were given 10-min to explore a clear plexiglass arena (45 × 45 × 30 cm) while they received photostimulation (10-Hz, 10-ms pulse width)*(21)*. The time spent in the center of the arena and ambulation were recorded using EthoVision v12-14.

#### Chemogenetic Inhibition of LC-NE Cell Bodies or LC-LHA Circuit During Anxiety & Feeding

##### Stress-induced anxiety in the open field test (OFT)

Mice received CNO (1-mg/kg i.p.) approximately 30-min before being placed into a restraint tube (TV-150 STD, Braintree Scientific) for 30-min, as described previously*(30)*. Immediately following restraint stress, mice were given 20-min to explore a novel open field arena (45 × 45 × 30 cm). The time spent in the center of the arena and total ambulation was recorded using EthoVision v15.

##### Food intake test (FI)

To motivate feeding, mice were overnight fasted prior to measuring food intake in a familiar arena. In LC^hM4Di^ cell body experiments, mice received CNO (1-mg/kg i.p.) or vehicle approximately 20-min before the FI test. For LC-LHA^hM4Di^ circuit experiments, mice were briefly anesthetized (4% isoflurane) to connect the internal cannulae to the guide cannulae and received bilateral local infusions of CNO (0.3-uM, 100 nL per side)*(76)* or aCSF (100-nL per side) at 100 nL/min through bilateral internal cannulae that projected 1.5 mm below the guide cannula (C235IS-5/SPC, P1 Technologies). Internal cannulae were left in place for 2-min to minimize backflow of the drug upon withdrawal. Mice were then returned to their homecages for 30-min until the FI test. Food intake was measured after 30-min by an experimenter blind to treatment group.

### Optogenetic Inhibition of LC-LHA Circuit During Feeding

LC-LHA^eNpHR^ mice and LC-LHA^EYFP^ controls were briefly anesthetized (4% isoflurane) to connect the fiber-optic cannula to bilateral polymer optical fibers (0.63 NA, 500-µm, 0.5-m length, Prizmatix), connected to an output port of a fiber optic rotary joint (Prizmatix). Mice were given 20-min to recover in the homecage before testing. During the food intake test (see *FI section* above for test details), mice received constant Lime-Green LED illumination (5-mW total power, Dual Channel Optogenetics-Lime-Green-LED module, Prizmatix). Food intake was measured for 30-min in free-feeding mice by an experimenter blind to treatment group.

### Statistical Analysis

ANOVAs and t-tests were used to determine differences between groups. Bonferroni posthoc tests were used as appropriate. Significance was set at *P*<0.05 for all analyses. All predictions were two-tailed unless otherwise stated to assess *a priori* predictions. All data are expressed as mean ± standard error (SEM). For consistency across experiments, we present food intake in grams (e.g., FI test) or number of pellets consumed (e.g., FED studies). Further, we report % change for experiments employing repeated measure(s). For quantification of Fos expression following LC^hM3Dq^ activation, we used one-tailed tests to assess a *priori* predictions. Tukey’s strategy (1.5 IQR above or below the 25^th^ or 75^th^ percentile) was used for detection and removal of extreme outliers to avoid a type II statistical error. This conservative strategy identified the following outliers that were excluded from analysis: an LC^mCherry^ and LC^hM4Di^ mouse (open field) and LC-LHA^ChR2^ mouse (in baseline food intake). Analyses were conducted using GraphPad Prism 7 or 8 (GraphPad Software Inc.) and IBM SPSS software for Windows, version 21 (SPSS Inc., Chicago IL). Statistics and sample size are listed in the figure legends.

## Author contributions

Conceptualization: NRS, PJ

Photometry: NRS, CMM under guidance of GC

Chemogenetics, Behavior and Metabolism: NRS, MH (experiments); MH, CAM, JA,

JMP, NRS (analysis) under guidance of AVK, MJK

Optogenetics and Behavior: NRS, MH under guidance of MRB, MJK, JDC

Stereotaxic Surgeries: NRS, LRW

Viral Constructs: NP

Immunohistochemistry, *In situ* hybridization, & Imaging: NRS, KGS, JA, CYK, JMP

Cell Counts: SAF, CXY

Supervision: NRS, PJ

Writing: NRS, PJ with input from coauthors

## Technical assistance

Behavior & Photometry Assistance: Philip Kindler, Kushal Prasad, and Juhee Haam Core Services: NIEHS Neurobehavioral, Fluorescence Microscopy & Imaging, and Viral Vector Cores

Animal Services: NIEHS Comparative Medicine Branch and Animal Resources

Contract Services: NIH, NIEHS Contract HHSN273201600011C with Social & Scientific Systems (Sandra McBride, Matthew Bridge, Mike Easterling) for photometry code.

## Data and materials availability

All data are available in the main text or the supplementary materials. The custom computer code (R, 64-bit version 3.5.1) used to align fluorescence signals was deposited to Github. Code can be accessed using the following link: https://github.com/NIEHS/FED-and-Photometry-Data-Processing.

## SUPPLEMENTARY MATERIALS

Supplementary Materials can be found with this article.

