## Supplementary Materials for "A Role for the Locus Coeruleus in the Modulation of Feeding"

**This PDF file includes:**

Supplementary Materials and Methods
Figs. S1 to S7
References (1 to 3)

### SUPPLEMENTARY MATERIALS AND METHODS

#### Immunohistochemistry

The antibodies used are summarized below.

| Antibody | Species | Dilution | Source | Catalog # | LOT # |
|---|---|---|---|---|---|
| mCherry | Rat | 1:1000 | Kerafast | EST202 | 105499, 032018 |
| EGFP | Chicken | 1:10,000 1:1000 | Abcam | AB13970 | 236651-15 |
| dsRed | Rabbit | 1:1000 | Clontech | 632496 | 1612022 |
| tdTomato | Rat | 1:2000 | Kerafast | EST203 | Not available |
| TH | Rabbit | 1:1000 | Millipore | AB152 | 2493925 |
| TH | Mouse | 1:500 | GeneTex | GTX10372 | 821901979, 821903616, 822101775 |
| c-Fos | Rabbit | 1:250 | Santa Cruz | Sc-52 | K1115 |
| c-Fos | Rabbit | 1:2000 | Abcam | ab190289 | GR289729-1 |
| Alexa Fluor 568 anti-rat | Goat | 1:1000 | Invitrogen | A-11077 | 1853640 |
| Alexa Fluor 488 anti-chicken | Goat | 1:1000 | Invitrogen | A-11039 | 1812246 |
| Alexa Fluor 488 anti-rabbit | Goat | 1:1000 | Invitrogen | A-11034 | 1812166 |
| Alexa Fluor 568 anti-rabbit | Goat | 1:1000 | Invitrogen | A-11036 | 1924788 |
| Alexa Fluor 633 anti-rabbit | Goat | 1:1000 | Invitrogen | A-21071 | 2199090 |
| Alexa Fluor 488 anti-mouse | Goat | 1:1000 | Invitrogen | A-11029 | 2306579 |
| Alexa Fluor 568 anti-mouse | Goat | 1:1000 | Invitrogen | A-11031 | 685228 |
| Alexa Fluor 633 anti-mouse | Goat | 1:1000 | Invitrogen | A-21052 | 1906490 |
| Biotinylated anti-rabbit | Goat | 1:500 | Vector Labs | BA-1000 | ZB0318 |

### Viral Preparation

The viruses used are summarized below.

| Plasmid | Virus source | Packaging | Serotype | Titer |
|---|---|---|---|---|
| pAAV-Ef1a-fDIO-GCaMP6f (addgene ID 128315) | Jensen lab (NIEHS) | NIEHS Viral Vector core | AAV5 | $1.05 \times 10^{13}$ vg $mL^{-1}$ |
| pAAV-Ef1a-fDIO-tdTomato (addgene ID 128434) | Jensen lab (NIEHS) | NIEHS Viral Vector core | AAV5 | $6.46 \times 10^{12}$ vg $mL^{-1}$<br>$1.05 \times 10^{13}$ vg $mL^{-1}$ |
| pAAV-EF1a-fDIO-ChrimsonR-tdTomato (addgene ID 171027) | Jensen lab (NIEHS) | NIEHS Viral Vector core | AAV5 | $1.65 \times 10^{13}$ vg $mL^{-1}$ |
| pAAV-hSyn-DIO-hM4D(Gi)-mCherry (addgene ID 44362) | Roth lab (UNC-Chapel Hill) | Addgene | AAV5 | $1.8 \times 10^{13}$ vg $mL^{-1}$ |
| pAAV-hSyn-DIO-mCherry (addgene ID 50459) | Roth lab (UNC-Chapel Hill) | Addgene | AAV5 | $8.4 \times 10^{12}$ vg $mL^{-1}$ |
| pAAV-Ef1a-DIO hChR2(H134R)-EYFP | Deisseroth lab (Stanford) | UNC Viral Vector core | AAV5 | $7 \times 10^{12}$ vg $mL^{-1}$ |
| pAAV-Ef1a-DIO EYFP | Deisseroth lab (Stanford) | UNC Viral Vector core | AAV5 | $4.4 \times 10^{12}$ vg $mL^{-1}$ |
| pAAV-Ef1a-DIO-eNpHR3.0-EYFP | Deisseroth lab (Stanford) | UNC Viral Vector core | AAV5 | $3.1 \times 10^{12}$ vg $mL^{-1}$ |

### Surgery

Surgery for Photometry Experiments. The locus coeruleus of *Dbh*$^{Flpo}$ mice were injected with a 1:5 viral cocktail of *Flp*-dependent GCaMP6f (AAV5-fEf1a-fDIO-GCaMP6f) and tdTomato (AAV5-Ef1a-fDIO-tdTomato) using the following stereotaxic coordinates: −5.45 posterior; +/- 1.0 lateral; -3.75 ventral (mm from bregma). For experiments combining chemogenetics with photometry, both triple transgenic *En1*$^{cre}$*; Dbh*$^{Flpo}$*; R/C::FL-hM3Dq* (LC$^{hM3Dq}$) and double transgenic *Dbh*$^{Flpo}$*; R/C::FL-hM3Dq* (LC$^{EGFP}$) mice were used. The locus coeruleus of LC$^{hM3Dq}$ mice were injected with a Flp-dependent GCaMP6f AAV, and a subset of these mice were also co-injected with a Flp-dependent tdTomato AAV at least 3 weeks before optical probe implantation. All mice were unilaterally implanted with optical probes above the LC (-5.45 posterior, ±0.85 lateral, -3.71 to -4.31 ventral, mm from bregma). Optical fiber probes were constructed by threading an optical fiber (200 µm, 0.39 NA, FT200EMT, Thorlabs; or 105 µm, 0.22 NA, Thorlabs) through a ceramic ferrule with a 1.25-mm OD (MM-CON2007–2300 or MM-CON2010-1270-2-WHT, Precision Fiber Products) and secured with heat-cured epoxy (Epoxy 353ND, Precision Fiber Products) and then polished and cleaved to length. Ventral placement of the optical fiber probe was guided by live fluorescence signal. During implantation of each probe, excitation light (470 nm, 760 mW mounted LED, M470L4, Thorlabs) was launched into a fluorescence cube (DFM1, Thorlabs) containing an excitation filter (ET470/40x, Chroma) and beam splitter (T495lpxt, Chroma). Excitation light was then reflected into an achromatic fiber port (PAF2-A4A, Thorlabs) and collected by a 5-meter-long relay multimode patch cable (M72L05, 200-µm core with 0.39 NA, Thorlabs) connected with a 1-meter-long multimode patch cable (M83L01: 200-µm core with 0.39 NA or M61L01: 105-µm core with 0.22 NA) linked to an implantable ferrule containing the optical fiber by a ceramic sleeve (SM-CS125S, Precision Fiber Products Inc.). Excitation light from the implantable fiber was adjusted to ~70-µW. Using the same optical fiber and patch cable, emitted light was passed through an emission filter (ET500lp, Chroma) before collection by a multi-mode patch cable. Emitted light was then collected into a spectrometer (Ocean FX, Ocean Insight) and visualized using Ocean View version 1.6.7. Optical probes were adhered with epoxy (PFP 353ND, Precision Fiber Products, Inc.) and anchored to the skull using 3 skull screws, Metabond (S371, S398, and S396, Parkell Inc.) and dental acrylic (1406R and 1230CLR, Lang Dental Manufacturing Co. Inc.).

Surgery for ChrimsonR Optogenetic Experiments. $Dbh^{Flpo}$ mice received bilateral LC injections of the Flp-dependent virus encoding ChrimsonR-tdTomato (AAV5-EF1a-fDIO-ChrimsonR-tdTomato) or tdTomato control (AAV5-EF1a-fDIO-tdTomato) using the following coordinates: -5.45 posterior, ±1.28 lateral, -3.65 mm ventral (mm from bregma)(*1*). Three weeks later, mice were implanted with bilateral fiber-optic cannula (200-µm, 0.39 NA, Thorlabs) angled 10° over LC using the following coordinates: -5.45 posterior, ±1.52 lateral, -3.50 to -3.65 ventral (mm from bregma) and angled 10° over the LHA using the following coordinates: -1.70 posterior, ±1.71 lateral, -4.81 to -4.86 ventral (mm from bregma). Ventral placement of the optical probe (200-µm, 0.39 NA, Thorlabs) was guided by live fluorescence signal, as described above in the *Surgery for Photometry Experiments* section. The implants were secured using Metabond and dental acrylic. We performed histology on every mouse to verify expression of virus and fiber placement. Importantly, we observed selective expression of ChrimsonR-tdTomato in TH+ locus coeruleus neurons, but no expression in other TH+ noradrenergic nuclei (e.g., A1, A2, A5) in all ChrimsonR mice.

Surgery for ChR2 and eNpHR Optogenetic Experiments. $Dbh^{cre}$ mice received bilateral LC injections of the cre-dependent virus encoding ChR2-EYFP (AAV5-Ef1a-DIO-ChR2-EYFP), eNpHR (AAV5-Ef1a-DIO-eNpHR3.0-EYFP), or EYFP control (AAV-Ef1a-DIO-EYFP) using the following coordinates: -5.45 posterior, ±1.28 lateral, -3.65 mm ventral (mm from bregma)(*1*). During the same surgery, mice were implanted with bilateral fiber-optic cannula (MFC_200/250-0.66_5mm_ZF1.25_FLT, Doric Lenses Inc., Quebec Canada) angled 10° over the LHA using the following coordinates: -1.70 posterior, ±1.71 lateral, -4.82 ventral (mm from bregma)(*2*). The implants were secured using Metabond and dental acrylic. We performed histology on every mouse to verify expression of virus and fiber placement. Mice with selective expression of ChR2 or eNpHR in TH+ locus coeruleus neurons, but no expression in other TH+ noradrenergic nuclei (e.g., A1, A2, A5) were included for analysis.

Surgery for Gi-DREADD Experiments. $Dbh^{cre}$ mice received bilateral LC injections of the cre-dependent virus encoding hM4Di (AAV5-hSyn-DIO-hM4Di-mCherry) or mCherry control (AAV5-hSyn-DIO-mCherry) using the following coordinates: -5.45 posterior, ±1.28 lateral, -

3.65 mm ventral (mm from bregma)(*1*). In the same surgery, mice were implanted with bilateral cannulae (C235GS-5-2.0/SPC, P1 Technologies, Roanoke VA) over the LHA using the following coordinates: -1.70 posterior, ±1.00 lateral, -3.45 ventral (mm from bregma)(*2*). Bilateral cannulae were secured using Metabond and dental acrylic. We performed histology on every mouse to verify expression of virus and cannula placement. Mice with selective bilateral expression of hM4Di in TH+ locus coeruleus neurons, but no expression in other TH+ noradrenergic nuclei (e.g., A1, A2, A5) were included for analysis.

**LC-NE Photometry Recordings**

Photometry Recordings for Sensory Experiments. A 1-s flash of light was presented every minute during the recording session which lasted ~30-min. The light source (Dual Channel Optogenetics-Lime-Green-LED module, Prizmatix, Givat-Shmuel, Israel) was connected to a polymer optical fiber (500-µm, NA 0.63, Prizmatix) that was positioned indirectly above the testing arena (Phenotyper).

Photometry Recordings for Feeding Experiments. Prior to recordings, mice were habituated for several days to eating from the feeding experimentation device (FED)(*3*) in both the homecage and testing arena (Phenotyper). The FED dispensed grain-based chow (25-mg pellets, Test Diet 5TUM) with a 60-s delay after retrieval of each pellet. This delay was used to space out pellets, which facilitated analysis of consumption responses during LC-NE calcium recordings in overnight fasted mice. Recordings collected with the FED lasted 60-min.

Photometry Recordings for Pharmacology Studies. Recordings lasted 35-min including 15-min prior to injection and 20-min post-injection. In all experiments, data were collected at 25-Hz from the camera and spectrometer (and FED when appropriate) and were synchronized using EthoVision (Noldus).

**Photometry Data Analysis**

Analysis of LC-NE Recordings During Feeding and Presentation of Light Flashes. We used RStudio (64-bit version 1.2.1335, RStudio, Inc., Boston, MA, USA) to run custom computer code (see *Code Availability* section) to align fluorescence ratio signals to relevant experimental events (i.e., pellet retrieval for feeding experiments, light flash for sensory experiments). The fluorescence ratio signal was assessed independently for each event around a 90-sec window for feeding experiments and 30-sec window for visual experiments. Each fluorescence ratio value was transformed into a z-score by subtracting the mean of a baseline period and dividing the SD of the baseline period. For feeding experiments, the 90-sec window was divided into four behavioral windows: baseline (45 to 30 sec before pellet retrieval), approach (1 to 0 sec before pellet retrieval), consumption (1 to 6 sec after pellet retrieval) and recovery (25 to 35 sec after pellet retrieval). Analysis periods were selected based on visual observation of behavior. To assess if LC activity varies as mice approached satiety, we compared LC responses during approach and consumption during the first and last ten pellets of the recording session.

For visual experiments, the 30-second window was divided into four behavioral windows: baseline (15 to 5 sec before light onset), pre-light (5 to 0 sec before light onset), light (0 to 1 sec), and post-light (5 to 15 sec after light onset). An average of the z-scores within the behavioral windows were reported and then averaged across events for each subject. Perievent rasters' and associated example traces were plotted using Neuroexplorer 5.109 (64-bit version). All other traces were plotted using GraphPad Prism 7 or 8 (GraphPad Software Inc., La Jolla CA).

Analysis of LC-NE Recordings During Chemogenetic Experiments. For quantification of CNO or vehicle effects on LC activity, fluorescence z-scores were calculated using the 5 minutes prior to the experimenter entering for injection to determine the baseline fluorescence mean and standard deviation. The mean fluorescence z-scores across 5-min bins were then calculated. The photometry data collected during scruffing and injection were excluded from quantification. These time points were manually marked in EthoVision v14 by an experimenter blind to treatment. Area under the curve of the fluorescence z-score traces during minutes 5-15 post injection were calculated in GraphPad Prism 8.

### SUPPLEMENTAL FIGURES

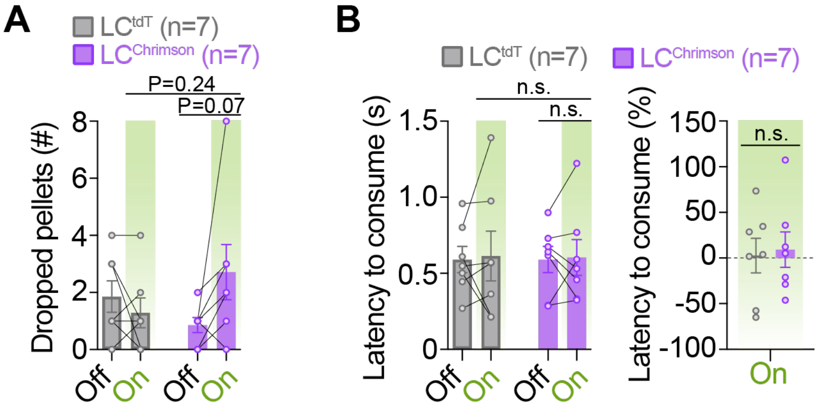


**Fig. S1. No significant change in rate of pellet drops or latency to consume during closed-loop optogenetic activation of LC-NE neurons. (A-B)** Feeding-related behaviors during the closed-loop optogenetic experiment wherein photostimulation (560-nm, 10-Hz, 10-ms pulses) was triggered upon pellet retrieval and occurred briefly for 10-sec during consumption from the feeding experimentation device (FED) in fasted mice. Two-way repeated measures ANOVA, stimulation x virus interaction: dropped pellets (**A.** $F_{1,12}$=4.898, $P$=0.0470) and average latency to consume (**B,** *Left*, $F_{1,12}$=0.004398, $P$=0.9482). Exact p-value shows the relevant Bonferroni post-hoc comparisons. n.s., non-significant. Unpaired samples *t*-test: percent latency to consume (**B.** *Right*, $t_{12}$=0.2406, $P$=0.8139). Data are mean ± SEM. $n$=7 LC$^{tdT}$ mice, $n$=7 LC$^{ChrimsonR}$ mice.

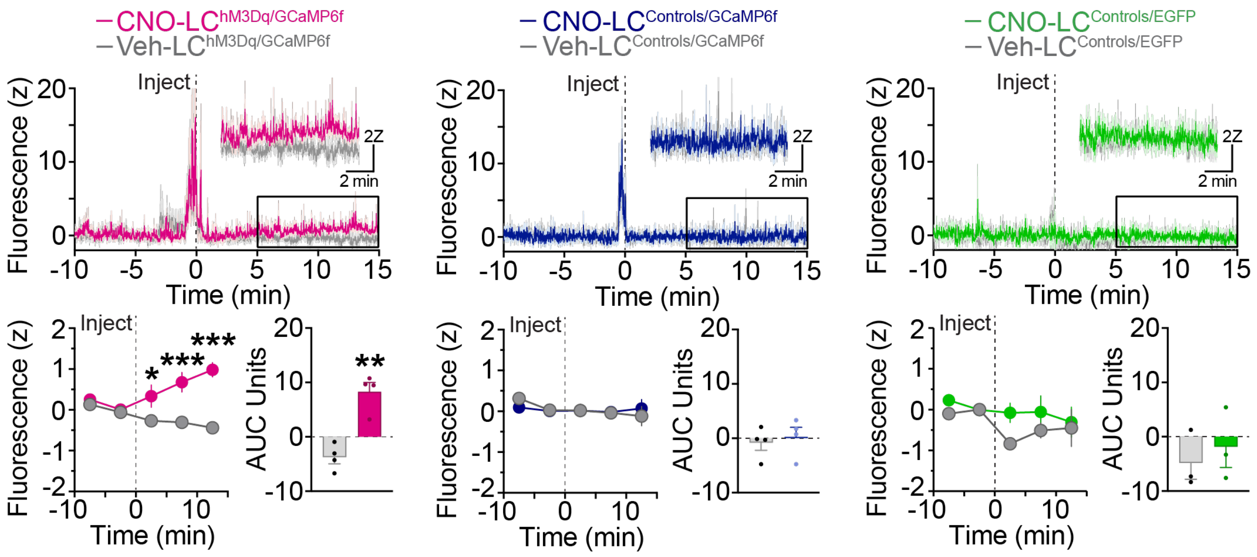


**Fig. S2. Selective chemogenetic activation of LC-NE neurons.** *Top.* Average z-score fluorescence ratio aligned to CNO (5 mg/kg i.p.) or vehicle injection in $LC^{hM3Dq/GCaMP6f}$ mice, littermate $LC^{Controls/GCaMP6f}$, and GCaMP6f-lacking littermate $LC^{Controls/EGFP}$. *Bottom left.* Average 5-min binned z-score fluorescence aligned to CNO or vehicle injection. Two-way repeated measures ANOVA, drug x time interaction: $LC^{hM3Dq/GCaMP6f}$ ($F_{4,12}$=11.99, $P$=0.0004), $LC^{Controls/GCaMP6f}$ ($F_{4,12}$=0.8213, $P$=0.5361), and $LC^{Controls/EGFP}$ ($F_{4,8}$=0.6928, $P$=0.6175). Bonferroni post-hoc test, ***$P$<0.01, *$P$<0.05 vs. vehicle. Data are mean ± SEM. $n$=4 $LC^{hM3Dq/GCaMP6f}$ mice, $n$=4 $LC^{Controls/GCaMP6f}$, and $n$=3 $LC^{Controls/EGFP}$. *Bottom right.* Area Under the Curve (AUC) from 5-min to 15-min following CNO or vehicle injection. Paired samples $t$-test: $LC^{hM3Dq/GCaMP6f}$ ($t_3$=10.22, **$P$<0.01), $LC^{Controls/GCaMP6f}$ ($t_3$=0.3990, $P$=0.7166), and $LC^{Controls/EGFP}$ ($t_2$=0.4699, $P$=0.6847). Data are mean ± SEM. $n$=4 $LC^{hM3Dq/GCaMP6f}$ mice, $n$=4 $LC^{Controls/GCaMP6f}$, and $n$=3 $LC^{Controls/EGFP}$.

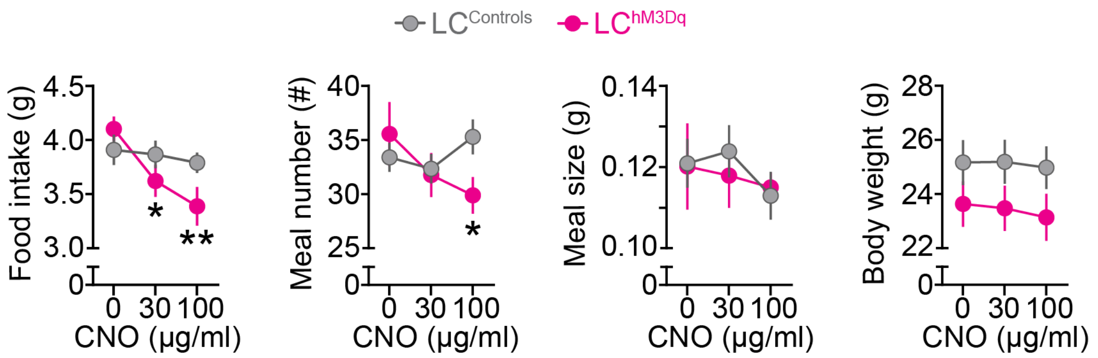


**Fig. S3. Chemogenetic activation of LC-NE neurons suppresses feeding.** Behavioral and metabolic measures in Labmaster automated homecage. Two-way repeated measures ANOVA, drug x genotype interaction: food intake ($F_{2,54}$=3.756, $P$=0.0297), meal number ($F_{2,54}$=4.518, $P$=0.0153), meal size ($F_{2,54}$=0.3750, $P$=0.6890), and body weight ($F_{2,70}$=2.559, $P$=0.0846). Bonferroni post-hoc test, **$P$<0.001, *$P$<0.01 vs. vehicle. Data are mean ± SEM. $n$=20 $LC^{Controls}$, $n$=9 $LC^{hM3Dq}$ mice for all measures except body weight wherein $n$=17 $LC^{hM3Dq}$ mice.

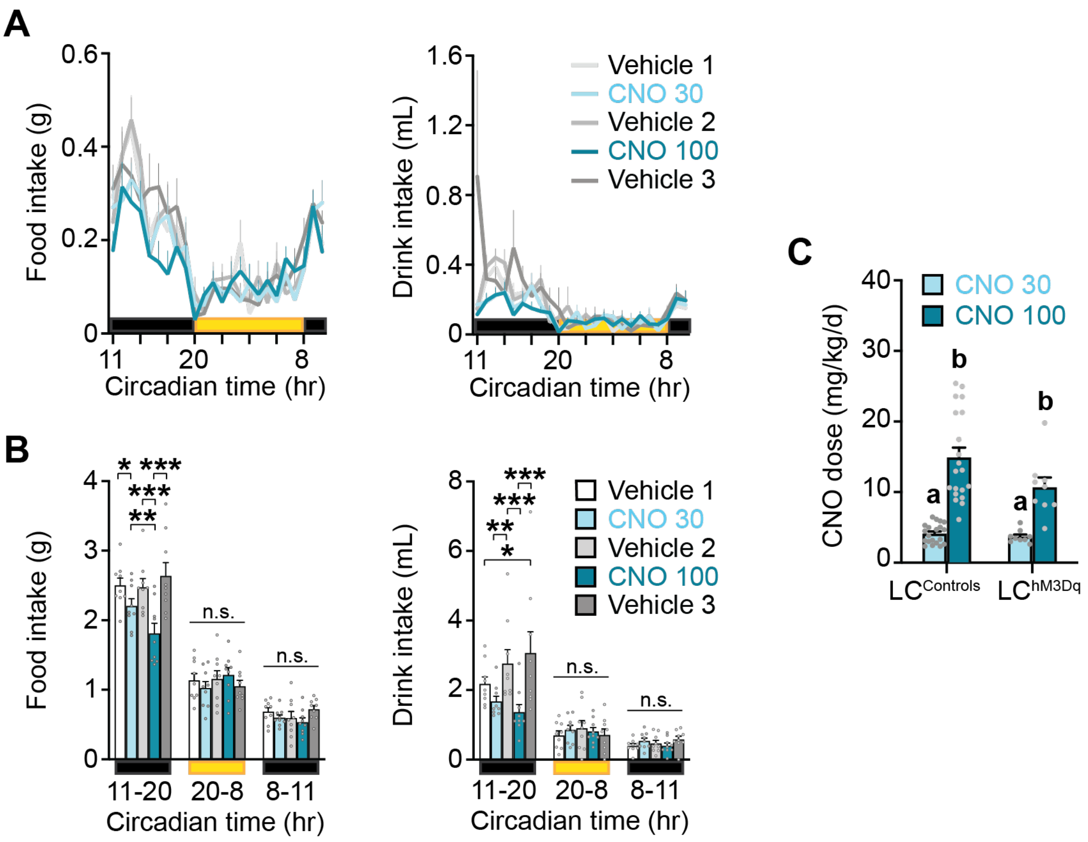


**Fig. S4. Chemogenetic activation of LC-NE neurons suppresses feeding and drinking during the early, dark-phase of the light cycle in a reversible manner. (A)** Circadian food and drink intake by LC$^{hM3Dq}$ mice receiving CNO (30 and 100 µg/mL) in drinking water. **(B)** Average food and drink intake during the 24-hour circadian light cycle. Two-way repeated measures ANOVA, drug x time interaction: food intake ($F_{8,64}$=8.02, $P$<0.001) and drink intake ($F_{8,64}$=3.93, $P$<0.001). Bonferroni post-hoc test, ***$P$<0.001, **$P$<0.01, *$P$<0.05. n.s., non-significant. Data are mean ± SEM. $n$=9 LC$^{hM3Dq}$ mice. CT, circadian time. Yellow shows circadian lights-on (CT 20-8). Black shows circadian lights-off, which was divided according to when drug was first introduced (CT 11-20; early dark) and later present (CT 8-11; late dark). **(C)** Average daily CNO dose by drinking water. Two-way repeated measures ANOVA, drug x genotype interaction: ($F_{1,27}$=3.995, $P$=0.0558). Main effect of drug: ($F_{1,27}$=87.96, $P$<0.0001), $P$<0.001 30 µg/mL (group a) vs. 100 µg/mL (group b). Data are mean ± SEM. $n$=20 LC$^{Controls}$, $n$=9 LC$^{hM3Dq}$ mice.

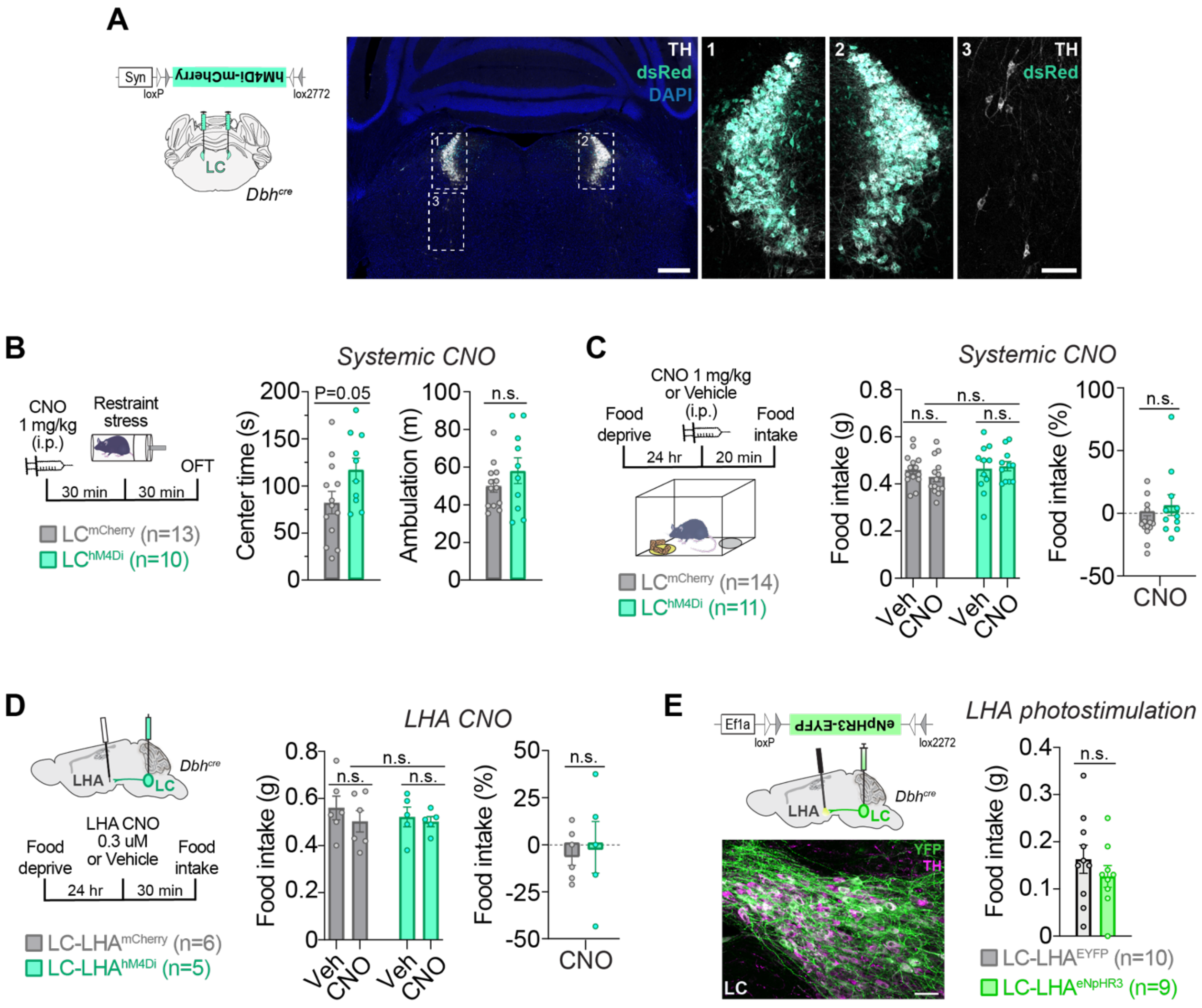


**Fig. S5. Inhibition of either LC-NE cell bodies or LC-LHA pathway results in no significant change in food intake. (A)** *Left.* Schematic illustration of a coronal mouse brain shows the location of cre-dependent AAV used to drive hM4Di-mCherry expression. *Right*. Immunofluorescent labeling shows expression of mCherry (dsRed, blue-green) in noradrenergic (TH, white) neurons of the bilateral LC (images 1 and 2) but not in noradrenergic ventral subcoeruleus (SubC, image 3) neurons in a coronal brain section from a LC$^{hM4Di}$ mouse. Scale is 400-µm (brain slice) and 100-µm (high-magnification images). **(B)** *Left.* Timeline of stress-induced anxiety experiments. *Right***.** Average time spent in the center of the open field test (OFT) and ambulation following systemic CNO delivery and stress exposure. Unpaired samples *t*-test: center time ($t_{21}$=2.036, *P*=0.0545) and ambulation ($t_{21}$=1.139, *P*=0.2674). Data are mean ± SEM. *n*=13 LC$^{mCherry}$ mice, *n*=10 LC$^{hM4Di}$ mice. **(C)** Food intake (FI) following systemic delivery of CNO. *Left.* Timeline of FI experiments. *Middle***.** Average FI in fasted mice in CNO and vehicle-treated mice. Two-way repeated measures ANOVA, drug x virus interaction: $F_{1,23}$=1.540, *P*=0.2271. n.s., non-significant. *Right.* CNO-evoked food intake as a percent change from vehicle. Unpaired samples *t*-test, $t_{23}$= 1.442, *P*=0.1629. Data are mean ± SEM. *n*=14 LC$^{mCherry}$ mice, *n*=11 LC$^{hM4Di}$ mice. **(D)** *Left.* Schematic illustration of sagittal mouse brain shows the location of cre-dependent AAV used to drive hM4Di-mCherry expression and drug infusion cannulae. Timeline of intra-LHA delivery of CNO (0.3 µM) before the FI assay. *Middle***.** Average food intake following intra-LHA delivery of CNO or vehicle in fasted mice. Two-way repeated measures ANOVA, drug x virus interaction: $F_{1,9}$=0.1953, *P*=0.6689. n.s., non-significant. *Right.* CNO-evoked feeding as a percent change from vehicle. Unpaired samples *t*-test, $t_9$= 0.2822, *P*=0.7842. Data are mean ± SEM. *n*=6 LC-LHA$^{mCherry}$ mice, *n*=5 LC-LHA$^{hM4Di}$ mice. **(E)** *Left.* Schematic illustration of a parasagittal mouse brain shows the location of cre-dependent AAV used to drive eNpHR3.0-EYFP expression and fiberoptic probes. Immunofluorescent labeling shows eNpHR3.0-EYFP expression (EYFP, green) in noradrenergic (TH, magenta) locus coeruleus neurons in a parasagittal brain section from a LC-LHA$^{eNpHR3}$ mouse. Scale, 50-µm. *Right.* Average food intake during photostimulation for 30-min in *ad libitum* fed mice. Unpaired samples *t*-test: $t_{17}$=0.9491, *P*=0.3559. Data are mean ± SEM. *n*=10 LC-LHA$^{EYFP}$ mice, *n*=9 LC-LHA$^{eNpHR3}$ mice.

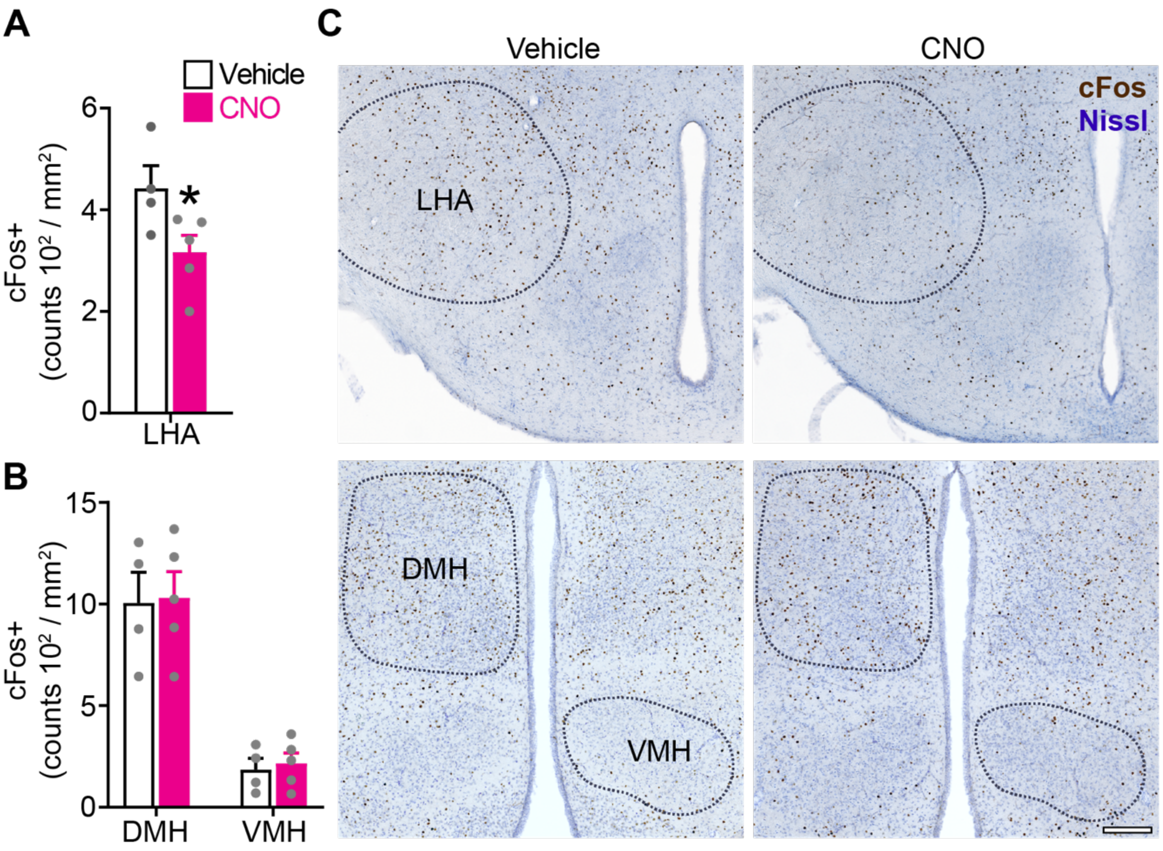


**Fig. S6. Chemogenetic activation of LC-NE neurons reduces neuronal activity in the LHA**. **(A-B)** Average Fos+ cell counts in the LHA of LC$^{hM3Dq}$ mice. Unpaired samples *t*-test (one-tailed): LHA (**A**, $t_7$=2.302, **P*<0.05), DMH (**B**, $t_7$=0.1281, *P*=0.4508), and VMH (**B**, $t_7$=0.3852, *P*=0.3558). Data are mean ± SEM. *n*=4 vehicle mice, *n*=5 CNO (1 mg/kg i.p.) mice. **(C)** Immunofluorescent labeling of coronal brain sections shows expression of Fos (brown) in hypothalamic nuclei (outlined) of LC$^{hM3Dq}$ mice. Tissue was counterstained with Nissl (blue). Scale, 100-µm. LHA, lateral hypothalamus area. DMH, dorsal medial hypothalamus. VMH, ventral medial hypothalamus.

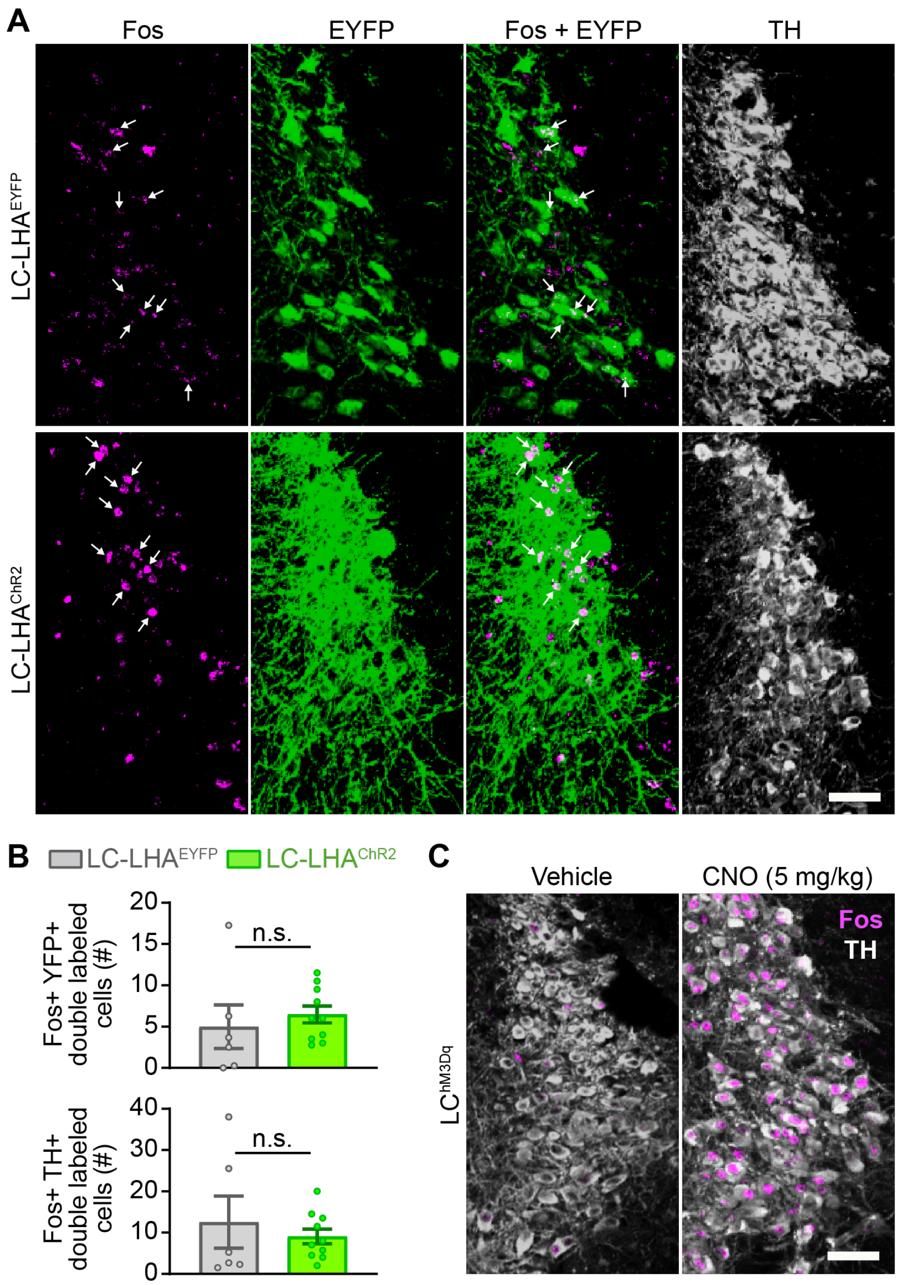


**Fig. S7. Stimulation of LC-LHA projection terminals does not influence LC-NE cell body activity. (A)** Immunofluorescent labeling of coronal brain sections shows co-expression of Fos (magenta) and EYFP (green) in noradrenergic (TH, white) locus coeruleus neurons of LC-LHA$^{ChR2}$ mice and LC-LHA$^{EYFP}$ controls. Arrows indicate location of double-labeled Fos+ YFP+ neurons in the locus coeruleus. Scale, 50-µm. **(B)** *Top.* Double-labeled Fos+ YFP+ neurons in the locus coeruleus. Unpaired samples *t*-test: $t_{14}$=0.6213, *P*=0.5444. *Bottom.* Double-labeled Fos+ TH+ neurons in the locus coeruleus. Unpaired samples *t*-test: $t_{14}$=0.6528, *P*=0.5244. Data are mean ± SEM. *n*=6 LC-LHA$^{EYFP}$ mice, *n*=10 LC-LHA$^{ChR2}$ mice. **(C)** Immunofluorescent labeling of coronal brain sections shows co-expression of Fos (magenta) in noradrenergic (TH, white) locus coeruleus neurons of vehicle-treated (negative control) and CNO-treated (positive control) LC$^{hM3Dq}$ mice, confirming our ability to detect Fos expression. Scale, 50-µm.
